# Glycosuria alters uropathogenic *Escherichia coli* global gene expression and virulence

**DOI:** 10.1101/2021.10.11.464023

**Authors:** Md. Jahirul Islam, Kamal Bagale, Preeti P John, Zachary Kurtz, Ritwij Kulkarni

**Affiliations:** Department of Biology, University of Louisiana at Lafayette, Lafayette, LA-70504

## Abstract

Uropathogenic *Escherichia coli* (UPEC) is the principal etiology of more than half of urinary tract infections (UTI) in humans with diabetes mellitus. Epidemiological data and studies in mouse model of ascending UTI have elucidated various host factors responsible for increasing the susceptibility of diabetic hosts to UPEC-UTI. In contrast, the nature of alterations in UPEC physiology mediated by diabetic urinary microenvironment and the contributions of altered UPEC physiology in shaping UPEC-UTI pathogenesis in diabetes have not been examined. Our central hypothesis is that glycosuria directly induces urinary virulence of UPEC. We compared virulence characteristics and gene expression in human UPEC strains UTI89 (cystitis) and CFT073 (pyelonephritis) exposed for 2h, *in vitro* to human urine either in the presence or absence of glycosuria (600mg/dl glucose). Compared to control UPEC exposed to nutrient-rich culture medium LB, glycosuria-exposed UPEC exhibited significant increase in biofilm formation and reduction in the hemagglutination of Guinea pig erythrocytes (a surrogate measure of type 1 piliation). In addition, analysis of UTI89 transcriptome by RNA sequencing revealed that 2h-long, *in vitro* exposure to glycosuria also significantly alters expression of virulence and metabolic genes central to urinary virulence of UPEC. In summary, our results provide novel insights into how glycosuria-mediated early changes in UPEC fitness may facilitate UTI pathogenesis in the diabetic urinary microenvironment.

**IMPORTANCE:** Uropathogenic *Escherichia coli* (UPEC) is an important causative agent of urinary tract infections in diabetic humans. We examined the effects of *in vitro* exposure to glycosuria (presence of glucose in urine) on the virulence and gene expression by UPEC. Our results show that glycosuria rapidly (in 2h) alters UPEC gene expression, induces biofilm formation, and suppresses hemagglutination. These results offer a novel insight into the pathogenesis of UPEC in the urinary tract.

## INTRODUCTION

Urinary tract infection (UTI) is a highly prevalent infection affecting an estimated 50% of women and 20% of men at least once during lifetime (1). Every year in the US, an estimated 10 million individuals with UTI symptoms seek medical help, of which around 400,000 require hospitalization (2, 3). UTI also complicates the prognosis of over a million hospital admissions every year(4). Gram negative uropathogenic *Escherichia coli* (UPEC) is the principal cause of >80% of community-acquired and > 40% hospital-associated UTI in humans (5). In addition, UPEC is also an important cause of more than half of all UTI cases in diabetic humans (6). The central premise of our research is that the increased susceptibility of diabetic individuals to UTI is a function of diabetes-mediated changes in not only host immune defenses but also the virulence of microbial pathogens. For this study, our major objective was to explore early changes in UPEC physiology caused by the short-term (2h-long) exposure to glycosuria. As model organisms we used two human UPEC strains: UTI89 isolated from a patient with cystitis and CFT073 isolated from a woman with pyelonephritis (7, 8). We exposed UPEC strains for 2h, *in vitro*, to plain human urine (denoted as U) or human urine supplemented with 600mg/dl glucose (UG) or to nutrient-rich lysogeny broth (LB) control. In this report, these exposures are presented as ‘name of the UPEC strain-exposure’; for example, UTI89-UG refers to UPEC strain UTI89 exposed for 2h to glycosuria.

Following exposure, we examined UTI89 gene expression using RNA sequencing (RNASeq) and quantitative real-time PCR (qRT-PCR). We also compared UTI89 and CFT073 exposed for 2h to LB, U, or UG using hemagglutination of Guinea pig RBCs (measure type 1 piliation) and in a mouse model of ascending UTI to identify changes in UPEC virulence. In addition, we examined biofilm formation by UTI89 and CFT073 in presence of LB, U, or UG. UTI89 qPCR and biofilm formation for UTI89 and CFT073 were also examined in urine supplemented with 600mg/dl galactose (U-Gal), to determine the contribution of osmolarity change on UPEC gene expression and biofilm formation.

Observational studies in humans and laboratory studies using experimental induction of ascending UTI in WT and diabetic mice have elucidated several host factors central to the increased susceptibility of diabetic individuals to UPEC UTI. For example, increased levels of advanced glycation end-products (AGE) in diabetic urinary tract are associated with bladder dysfunction; accumulation of AGE on urothelium facilitates urinary colonization by UPEC (9–11); diabetes-induced alterations in immune responses such as reduced antimicrobial peptides, pro-inflammatory cytokines, complement activation, and urinary leukocyte infiltration increase host susceptibility to UTI (12–16). On the pathogen side, glycosuria is shown to increase UTI risk by promoting growth of bacteria (17). However, whether glycosuria promotes bacterial virulence and whether a switch in bacterial metabolism toward energetically viable glycolytic pathways improves bacterial fitness in the diabetic urinary tract has not been deciphered. In this regard, we recently reported that glycosuria induces virulence of Gram positive *Streptococcus agalactiae* wherein *in vitro* exposure to human urine supplemented with glucose promoted bacterial (*S. agalactiae*) adherence to human bladder epithelium, resistance to antimicrobial peptide LL-37, and hemolytic ability (18). In this report we extend these findings and show that glycosuria significantly alters global transcriptome of UTI89 and induces virulence characteristics of UTI89 and CFT073. Overall, these findings further strengthen our central premise that glycosuria-mediated changes in the physiology of uropathogens plays a critical role in shaping pathogenesis of UTI in diabetics. We also note an important caveat that our experimental design does not examine how UPEC physiology may be affected by long-term exposure to glycosuria or by *in vivo* diabetic urinary microenvironment.

## MATERIALS AND METHODS

### Human urine collection

As approved by institutional review board at UL Lafayette, urine was collected from healthy female volunteers (18-45 years of age), who had not suffered from UTI and/or treated with antibiotics in a month prior to urine collection. Urine was sterilized using 0.22µm syringe filter and stored at −80°C in 1.5ml aliquots. On the day of experimentation, urine aliquots from at least 3 separate donors were pooled for use.

### Bacteria Strain and Growth Conditions

As model UPEC strains, we used UTI89, a bladder isolate from an acute cystitis patient and CFT073 a blood and urine isolate from a pyelonephritis patient (7, 8). Prior to experimentation, UPEC were cultured at 37°C in LB in a static culture for 24h followed by 1:100 subculture into fresh LB and incubation at 37°C, static for additional 24h. This cultivation protocol induces type 1 piliation in UPEC inoculum used for initiating ascending UTI in a murine model (19). In this report, UPEC cultures prepared in this manner are described as ‘type 1 piliated’.

Type 1 piliated UTI89 and CFT073 were pelleted, washed in sterile PBS, and then resuspended in LB, or in human urine (pooled from at least 3 donors) either plain (U) or supplemented with 600mg/dl glucose (UG) to mimic severe glycosuria or supplemented with 600mg/dl galactose (UGal) to cause equivalent change in osmolarity as UG. UPEC cultures were maintained at 37°C, static for varying lengths of time as indicated below.

### Growth curve

UPEC strains CFT073 and UTI89 were maintained 37°C in LB, U, or UG. CFUs were enumerated by dilution plating at 0, 1, 2, 4, 6, 8 and 24h time points. For calculating doubling time (DT), CFU/ml at 2h and 4h were used in the following formula:

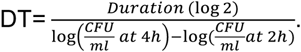

### In-vitro biofilm assay

Type 1 piliated UTI89 or CFT073 were pelleted, washed, and resuspended (1:100 dilution) in LB, U, UG, or UGal. Next, 100µl of each culture was added to the wells of a 96-well sterile polystyrene plate and incubated at 37°C and shaking at 150 rpm. We have previously described detailed protocols for biofilm visualization by crystal violet staining or CFU enumeration (20, 21). In short, after 3 rigorous washes with sterile PBS, biofilm-bound bacteria were scraped off with a pipette tip and enumerated by dilution plating. Planktonic CFU were determined by dilution plating the supernatant collected prior to washing. In separate experiments, rigorously washed biofilms were dried, baked at 60°C, and stained with crystal violet. Biofilm biomass is directly proportional to the retained crystal violet, which was extracted in ethanol-isopropanol mix and color intensity of the extract was measured at 590nm absorbance. To examine effects of iron-deprivation on biofilm formation, we resuspended type 1 piliated UPEC in LB, U, or UG containing 300μM DIP (2’,2’-dipyridyl), a membrane-permeable iron chelator or equal volume of DMSO (control-carrier for DIP) (22, 23). These biofilms were processed for either CFU enumeration or crystal violet staining.

### Hemagglutination of guinea pig RBCs

Type 1 piliated UTI89 and CFT073 were exposed to LB, U, or UG for 2h at 37°C, static. After exposure, bacterial pellets (5000rpm, 10min, 4°C) were resuspended in sterile PBS to obtain OD_540_=1.0 and maintained on ice. Citrated guinea pig blood (Colorado Serum Company) was centrifuged at 1,200 rpm, 4°C. RBC pellet was washed in cold sterile PBS until supernatant is clear (no red color). RBCs were set to OD_640_=1.9. For hemagglutination assay, 25µl bacteria were two-fold diluted in sterile PBS, 12 times (from 2^0^—undiluted to 2^11^) in three technical replicates in a V-bottom plate. 25µl blood was added to each well, mixed by tapping plate sides, covered with plate sealer, and incubated at 4°C overnight. For determining HA titer, plates were visually examined for the highest dilution showing RBCs as a red button at the bottom of the V-shaped well. Control wells contained LB, plain urine, or urine supplemented with glucose.

### RNA extraction, Reverse transcription, and Quantitative real-time PCR (qPCR)

Type 1 piliated UTI89 were pelleted, resuspended in LB, U, UG, or UGal and incubated for 2 h at 37°C, static. Bacterial RNA was extracted using Ambion® Ribopure^TM^ kit (Thermo Fisher Scientific) and treated with DNase. The quality (A260/A280) and quantity of extracted RNA was determined by nanodrop (Synergy™ HTX Multi-Mode Microplate Reader, Biotek). Next, 1µg total RNA was reverse transcribed using the Applied Biosystems^TM^ High-Capacity cDNA Reverse Transcription kit, (Thermo Fisher Scientific). Quantitative real-time PCR (qRT-PCR) was carried out using *Power* SYBR^®^ green master mix (Applied Biosystems) in StepOne™ Plus thermal cycler (Applied Biosystems) and relative quantification (RQ) of gene expression was determined using ΔΔC_T_ algorithm. List of primers used for qRT-PCR is shown in table 6.

**Table 6:**
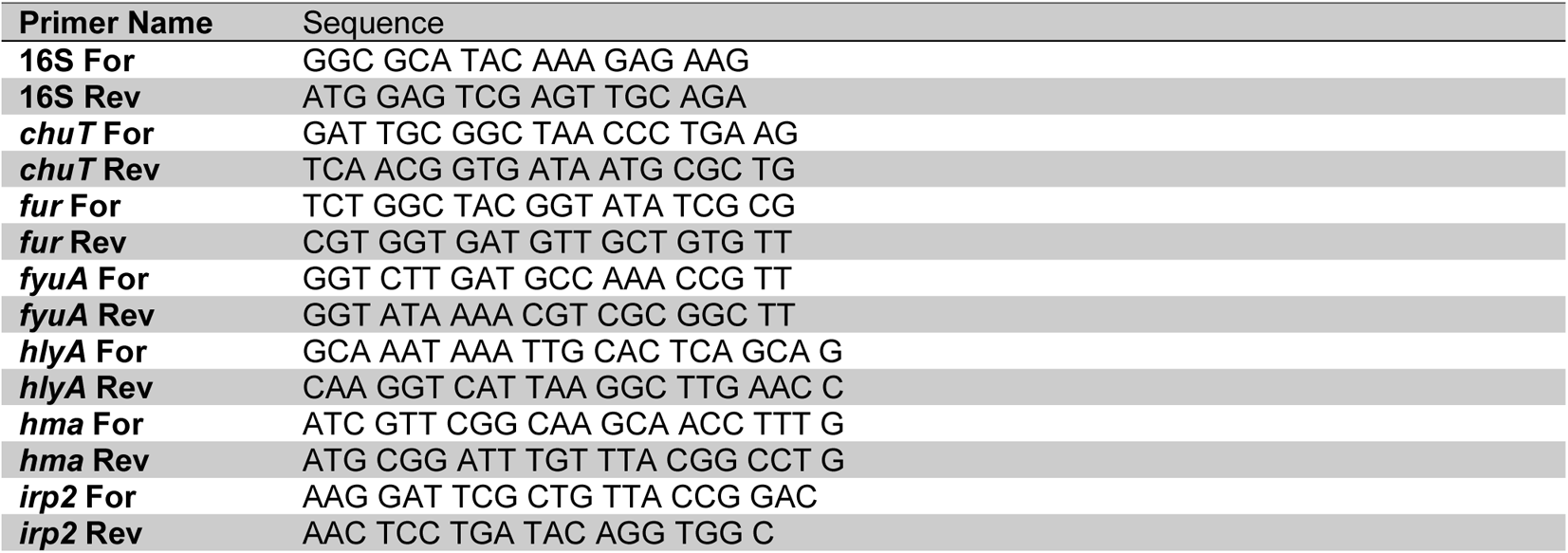
Primers used in this study

### Mouse model of ascending UTI

To examine changes in the virulence of UPEC mediated by exposure to glycosuria, male and female C3H mice were infected via transurethral catheterization with 10^7^ CFUs of UTI89 or CFT073 pre-exposed to LB, U, or UG (2h, 37°C, static). The protocol for transurethral catheterization of uropathogens to induce ascending UTI is previously described in details (19). At 24h post-infection (hpi), bacterial burden in bladder, kidneys, and spleen was determined by dilution plating the organ homogenates.

### Statistical analysis

Data from multiple biological replicates for each experiment were pooled together. Graphing and statistical analyses were done using GraphPad Prism 9.2.0. Results are expressed as the means ± standard deviation from data collected from two or more biological replicates each with two or more technical replicates. The data were compared by one-way ANOVA followed by Tukey’s multiple comparisons or by Student’s t test. Organ burden data are presented as scatter plots with median as central tendency. These were compared using Mann-Whitney U statistic. The difference between groups is considered significant if *P* ≤ 0.05.

### RNA-seq and data analysis

For RNA-Seq, total RNA was extracted from UTI89 exposed to LB, U, or UG (2h, 37°C, static) as described above. All library construction and initial analysis of differential expression was done by GENEWIZ (New Jersey, USA). Library construction included DNase treatment (TURBO™ DNase, ThermoFisher Scientific) and rDNA depletion (QIAseq FastSelect, Qiagen) followed by RNA fragmentation and random priming. cDNA synthesis (NEBNext® Ultra™ II, New England Biolabs) was followed by end repair, 5’ phosphorylation and dA-tailing. Libraries were sequenced on a partial lane of Illumina HiSeq 4000 with 150bp PE sequencing. Quality of sequence data was assessed using FastQC. All reads were quality filtered and trimmed using Trimmomatic v 0.36 with default settings (24). Reads were mapped to *Escherichia coli* O18:K1:H7 UTI89 (UPEC) genome using bowtie v 2.2.6 (25) and hit count for individual genes were generated using the featurecounts command in the subreads package v 1.5.2 (26). Genes with <10 reads were dropped from the analysis for differential expression. Differential expression for each gene was assessed using Wald tests implemented in DESeq2 (27). Genes with an adjusted p-value < 0.05 and absolute log_2_ fold change > 1 were categorized as differentially expressed genes (DEG).

GO enrichment analyses were conducted using the *goseq* v1.42.0 package in R (28). Gene ontology (GO) terms and gene lengths were extracted for each gene in *Escherichia coli* O18:K1:H7 UTI89 (UPEC) from the UniProt website (29). We determined whether DEGs between treatments were significantly overrepresented within molecular function, biological process, and cellular component GO terms using a Wallenius approximation and accounting for gene length bias using a probability weight function. Because of the inherent difficulties with multiple testing and correcting for multiple testing in GO analyses, we simply consider any term with p < 0.01 as statistically significant. KEGG pathway enrichment analyses were conducted using the KEGGREST Bioconductor package v1.30.1 (30) and a custom script (available upon request). KEGGREST was used to download lists of pathways and genes within pathways from the KEGG website for *Escherichia coli* O18:K1:H7 UTI89 (UPEC) (organism code *eci*).

We assessed whether DEGs are overrepresented in certain pathways using a Wilcoxon rank-sum test to determine whether adjusted p-values for differential expression of genes within a focal pathway are less than the adjusted p-values for differential expression of genes that are not within the pathway.

## RESULTS

### UTI89 and CFT073 grow in human urine both in the presence or absence of glucose

To confirm that UPEC survive in human urine either in the presence or absence of glucose, we cultivated type 1 piliated UTI89 and CFT073 in LB (medium control), U, or UG at 37°C for 24h under static conditions. CFUs were enumerated at 0, 1, 2, 4, 6, 8 and 24h. We did not observe statistically significant change in the doubling times of either UTI89-U, UTI89-UG or CFT073-U, CFT073-UG in comparison to their counterparts in LB (Fig 1 A, C). Although compared to their respective LB controls, both UTI89 and CFT073 cultivated in either U or UG showed statistically insignificant, 5-8-fold reduction in CFU/ml at 24h time point (Fig 1B, D).

**Figure 1:**
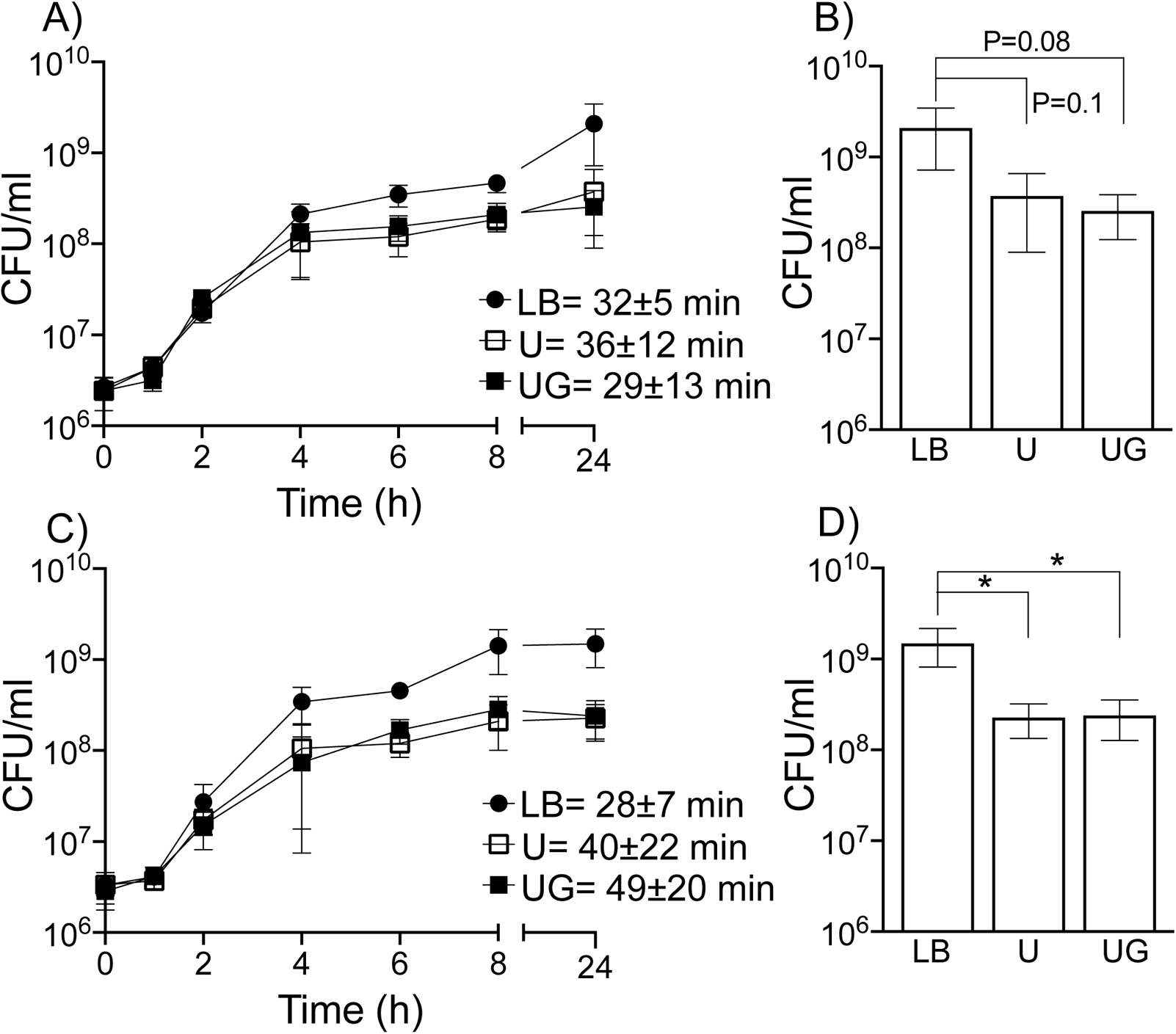
Growth of UPEC strains in presence of LB, plain urine, or glycosuria. We monitored growth of UTI89 (A, B) or CFT073 (C, D) in LB (control), human urine plain (U), or in urine supplemented with 600mg/dl glucose by enumerating CFU/ml at various time points up to 24h. For every time point average CFU/ml (from a minimum of two biological replicates) ± standard deviation is shown (A, C). Average doubling time (in minutes) ± standard deviation is shown next to the plot for each specific exposure (A, C). CFU/ml at 24h are shown separately (B, D) to facilitate comparison between final bacterial counts in each exposure.

### RNA-Seq and read mapping

We used RNA-Seq to quantify differential gene expression by assessing variation across the transcriptomes of UTI89-LB (control), UTI89-U, UTI89-UG. UTI89 was selected for RNA-Seq as it is a well-studied bladder isolate from acute cystitis patient (7). RNA was isolated from three independent biological replicates in each treatment. Whole transcriptome sequencing with rRNA depletion resulted in an average of 23 million 150bp paired-end reads per sample (range: 14-37.4 million reads). After adapter trimming and quality filtering, we retained an average of 96.5% of reads (94.7-98.6%) per library.

Subsequently, an average of 21.7 million reads (13.4 to 34.7 million reads) per library was successfully mapped to *Escherichia coli* UTI89 (O8:K1:H7) reference genome (GenBank accession no. CP000243). Euclidean distances between samples and principal component analysis for comparisons between UTI89-LB versus UTI89-U (Fig S1), UTI89-LB versus UTI89-UG (Fig S2), and UTI89-U versus UTI89-UG (Fig S3), revealed that expression patterns in biological replicates of each treatment group were more similar to each other than they were to those in biological replicates of the contrasting treatment group. Detailed RNA-Seq data (normalized counts for triplicate samples), log_2_FC (fold change), and Padj (adjusted P value) for all genes are presented in supplementary tables S1 (UTI89-LB versus UTI89-U), S4 (UTI89-UG versus UTI89-LB), and S7 (UTI89-U versus UTI89-U). Within UTI89-LB versus UTI89-U comparison, 638 genes showed significantly altered expression (defined as |log2FC| >1 and Padj ≤ 0.05), of which 393 were significantly upregulated (Supplementary Table S2), while and 245 were significantly downregulated (Supplementary Table S3). Within UTI89-LB versus UTI89-UG comparison, of 846 differentially expressed genes, 395 were significantly upregulated (Supplementary Table S5) and 245 significantly downregulated (Supplementary Table S6). Within UTI89-UG versus UTI89-U comparison, of the 691 differentially expressed genes, of which 253 were significantly upregulated (Supplementary Table S8) and 438 significantly downregulated (Supplementary Table S9). Fig 2 shows Venn diagrams showing differentially expressed genes (Fig 2A) and up-/down-regulated genes (Fig 2B) from different exposures. RNASeq results were confirmed by qRTPCR. Fold change for LB versus U and LB versus UG was correlated across the genes with r^2^=0.83 and r^2^=0.93, respectively (supplementary Fig S4).

**Figure 2:**
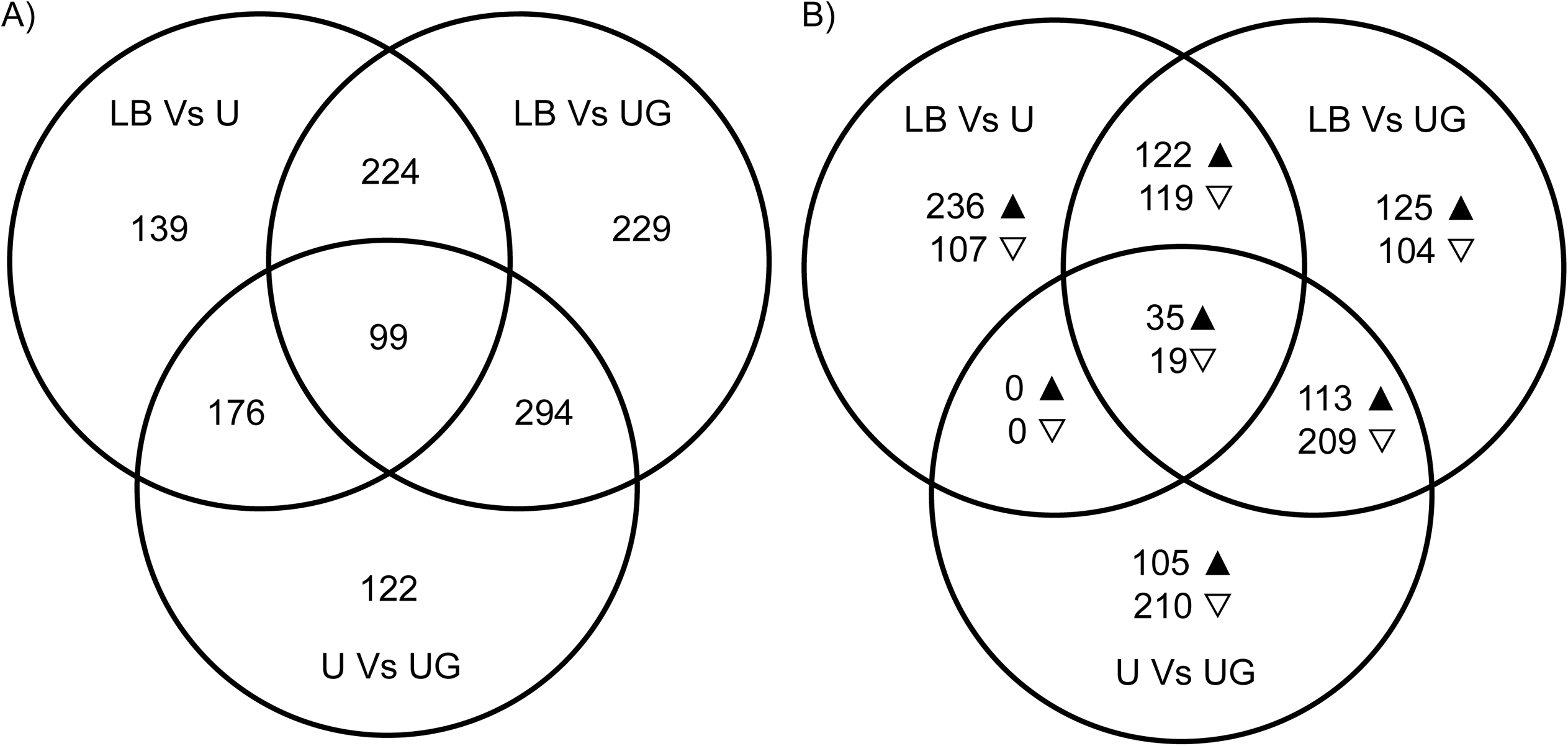
Comparison of differentially expressed genes (DEG) among different exposures. A) The total number of statistically significantly DEG and B) the numbers of significantly upregulated (p) and downregulated (s) genes are shown for the three comparisons, UTI89-U versus UTI89-LB, UTI89-UG versus UTI89-LB, and UTI89-UG versus UTI89-U. The results represent analysis of RNA-Seq data from three independent biological replicates. Differential expression was assessed using a threshold of absolute log_2_FC*≥*1 and Padj*≤*0.05.

**Figure 3:**
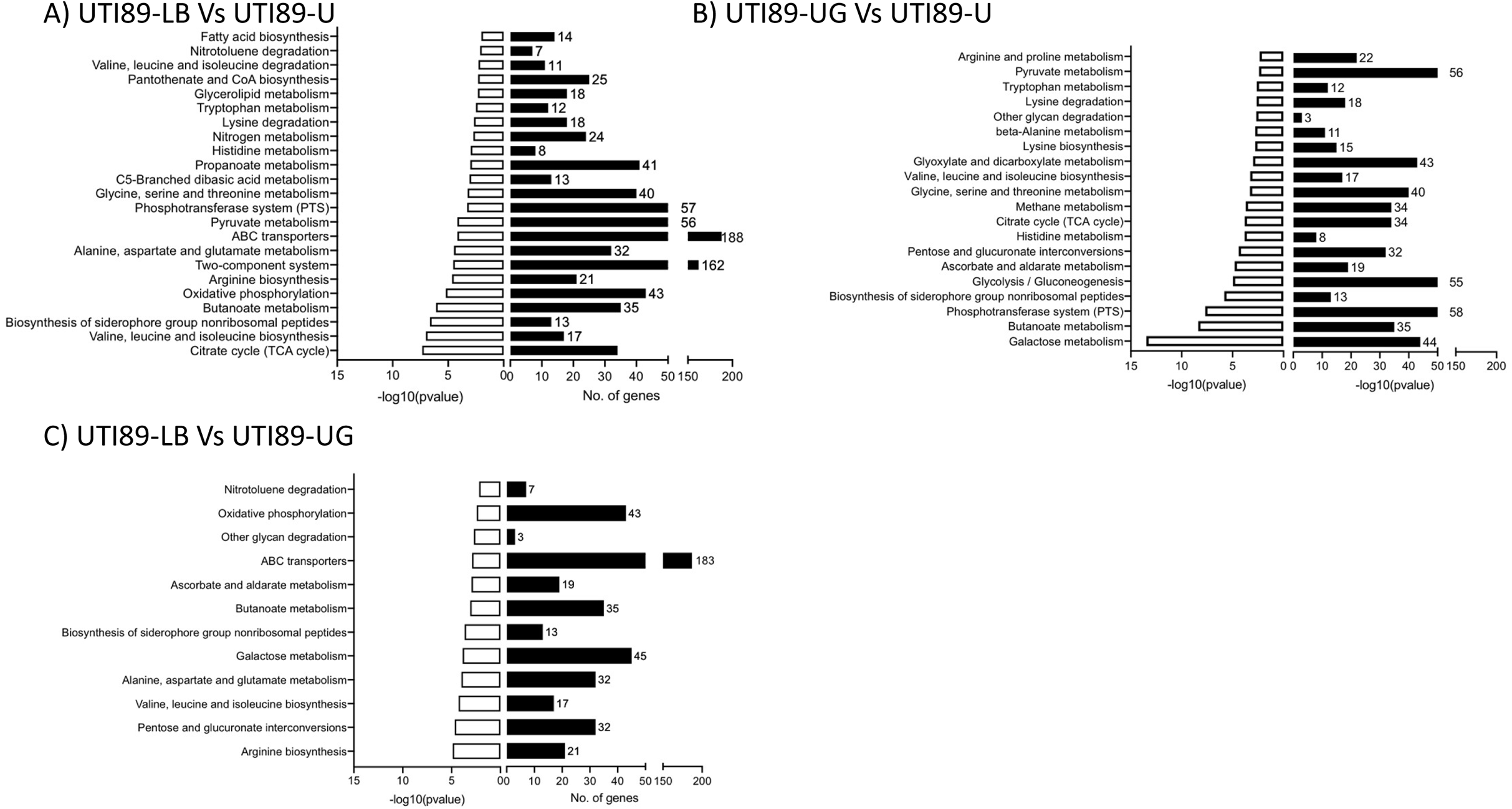
Enrichment analysis of RNA-Seq data. KEGG pathways significantly enriched for DEG at *P*<0.01 are shown for three comparisons (A) UTI89-U versus UTI89-LB, (B) UTI89-UG versus UTI89-LB, and (C) UTI89-UG versus UTI89-U. For each KEGG pathway, the number of DEG is shown inside the histogram while the total number of annotated genes is shown next to the histogram.

### Glycosuria affects expression of UTI89 genes involved in urinary fitness

The lower gastro-intestinal tract is a primary and immediate reservoir for UPEC before it gains access to the urinary tract via fecal contamination of urethra and causes ascending UTI (31, 32). When transitioning from nutritionally rich gastro-intestinal microenvironment to the healthy (non-diabetic) urinary tract, UPEC switches to catabolizing amino acids and short peptides present in low concentration in urine using gluconeogenesis; in individuals with glycosuria, however, glucose in urine is expected to serve as a ready source of carbon that is utilized via glycolysis (33, 34). In response to iron-limiting urinary microenvironment, UPEC also show upregulation iron acquisition system genes (35). Hence, we examined UTI89 transcriptomes for the differential expression of genes encoding specific metabolic and regulatory involved in peptide transport, glucose metabolism, and metal acquisition systems, which are defined as critical determinants of metabolic fitness of UPEC in the urinary tract (Tables 1–3). Given our primary interest in UTI pathogenesis, we also examined expression of genes encoding UPEC virulence factors (adhesins, toxins, regulators, and genes from KEGG pathway *eci02026* titled biofilm formation, Table 4) and genes categorized under human diseases (Table 5). In tables 1–5, differential gene expression data for UTI89-LB versus UTI89-U, UTI89-LB versus UTI89-UG, or for UTI89-U versus UTI89-UG comparisons is presented only where |log_2_FC|<1 and *P_adj_* <0.05.

**Table 1:** RNA-Seq analysis of expression of genes involved in amino acid and peptide transport

| KEGG ID | Gene symbol | LB vs U<br>log<sub>2</sub>FC<sup>a</sup> | LB vs UG<br>log<sub>2</sub>FC<sup>a</sup> | U vs UG<br>log<sub>2</sub>FC<sup>a</sup> | GenBank definition |
| --- | --- | --- | --- | --- | --- |
| UTI89_C0213 | *metQ* | -0.5782 | 2.9934* | 3.5851* | methionine ABC transporter substrate-binding lipoprotein MetQ |
| UTI89_C0215 | *metN* | -0.6530 | 3.5251* | 4.1934* | methionine ABC transporter ATP-binding protein MetN |
| UTI89_C0650 | *gltJ* | 1.3098* | 0.8220 | -0.4727 | glutamate/aspartate ABC transporter permease GltJ |
| UTI89_C0651 | *gltI* | 1.8123* | 1.4735* | -0.3174 | glutamate/aspartate ABC transporter substrate-binding protein GltI |
| UTI89_C0812 | *glnQ* | 1.1794* | 1.2137* | 0.0517 | glutamine ABC transporter ATP-binding protein GlnQ |
| UTI89_C0813 | *glnP* | 1.4902* | 1.4623* | -0.0094 | glutamine ABC transporter permease GlnP |
| UTI89_C0814 | *glnH* | 1.4642* | 1.7125* | 0.2684 | glutamine ABC transporter substrate-binding protein GlnH |
| UTI89_C0863 | *artJ* | 5.9098* | 6.8334* | 0.9428 | ABC transporter substrate-binding protein ArtJ |
| UTI89_C0864 | *artM* | 1.5315 | 1.4026* | -0.1162 | arginine ABC transporter permease ArtM |
| UTI89_C0865 | *artQ* | 1.4939 | 1.3493* | -0.1311 | arginine ABC transporter permease ArtQ |
| UTI89_C0866 | *artJ* | 1.9414* | 1.7229* | -0.2028 | arginine ABC transporter substrate-binding protein |
| UTI89_C0867 | *artP* | 1.4995* | 1.4418* | -0.0409 | arginine ABC transporter ATP-binding protein ArtP |
| UTI89_C1601 | *mppA* | 0.1456 | -1.3388* | -1.4674* | murein tripeptide ABC transporter substrate-binding protein MppA |
| UTI89_C2403 | *yehY* | -0.9065 | -1.3267* | -0.4080 | glycine betaine ABC transporter permease YehY |
| UTI89_C2592 | *hisQ* | 1.8122* | 1.7390* | -0.0528 | histidine ABC transporter permease HisQ |
| UTI89_C2593 | *hisJ* | 1.4887* | 1.5513 | 0.0823 | histidine ABC transporter substrate-binding protein HisJ |
| UTI89_C3037 | *proV* | -2.5853* | -3.2195* | -0.6216 | glycine betaine/L-proline ABC transporter ATP-binding protein ProV |
| UTI89_C3038 | *proW* | -2.7788 | -2.9565* | -0.1639 | glycine betaine/L-proline ABC transporter permease ProW |
| UTI89_C3039 | *proX* | -1.5426* | -1.6927* | -0.1348 | glycine betaine/L-proline ABC transporter substrate-binding protein ProX |
| UTI89_C3527 | *sstT* | 1.7573* | 1.0781* | -0.6618 | serine/threonine transporter SstT |
| UTI89_C3961 | *livF* | 1.1115 | 1.4958* | 0.3984 | high-affinity branched-chain amino acid ABC transporter ATP-binding protein LivF |
| UTI89_C3962 | *livG* | 1.2731* | 1.7045* | 0.4480 | high-affinity branched-chain amino acid ABC transporter ATP-binding protein LivG |
| UTI89_C3963 | *livM* | 1.3353* | 1.6741* | 0.3534 | branched chain amino acid ABC transporter permease LivM |
| UTI89_C3964 | *livH* | 2.1604* | 1.8579* | -0.2888 | high-affinity branched-chain amino acid ABC transporter permease LivH |
| UTI89_C3965 | *livK* | 2.8903* | 2.8508* | -0.0264 | high-affinity branched-chain amino acid ABC transporter substrate-binding protein LivK |
| UTI89_C3969 | *livJ* | 1.6578* | 1.6359* | -0.0044 | branched chain amino acid ABC transporter substrate-binding protein LivJ |
| UTI89_C4081 | *dppB* | 1.9892* | 0.8777 | -1.0933 | dipeptide ABC transporter permease DppB |
| UTI89_C4082 | *dppA* | 2.2105* | 1.1485* | -1.0444 | dipeptide ABC transporter substrate-binding protein DppA |
| UTI89_C4817 | *cycA* | 1.1040* | 0.5181 | -0.5676 | D-serine/D-alanine/glycine transporter |
| UTI89_C4956 | *dsdX* | 1.2850* | 0.5240 | -0.7400 | D-serine transporter DsdX |

<sup>a</sup> Only values where absolute log<sub>2</sub>[fold change] >1 and P<sub>adj</sub><0.05 (denoted by \*) in at least one of the three comparisons are shown. G<sup>b</sup> refers to glycolysis, N<sup>c</sup> to gluconeogenesis, and TCA<sup>d</sup> to TCA cycle. <sup>e</sup> Genes encode enzymes catalyzing pyruvate oxidation.

**Table 2:**
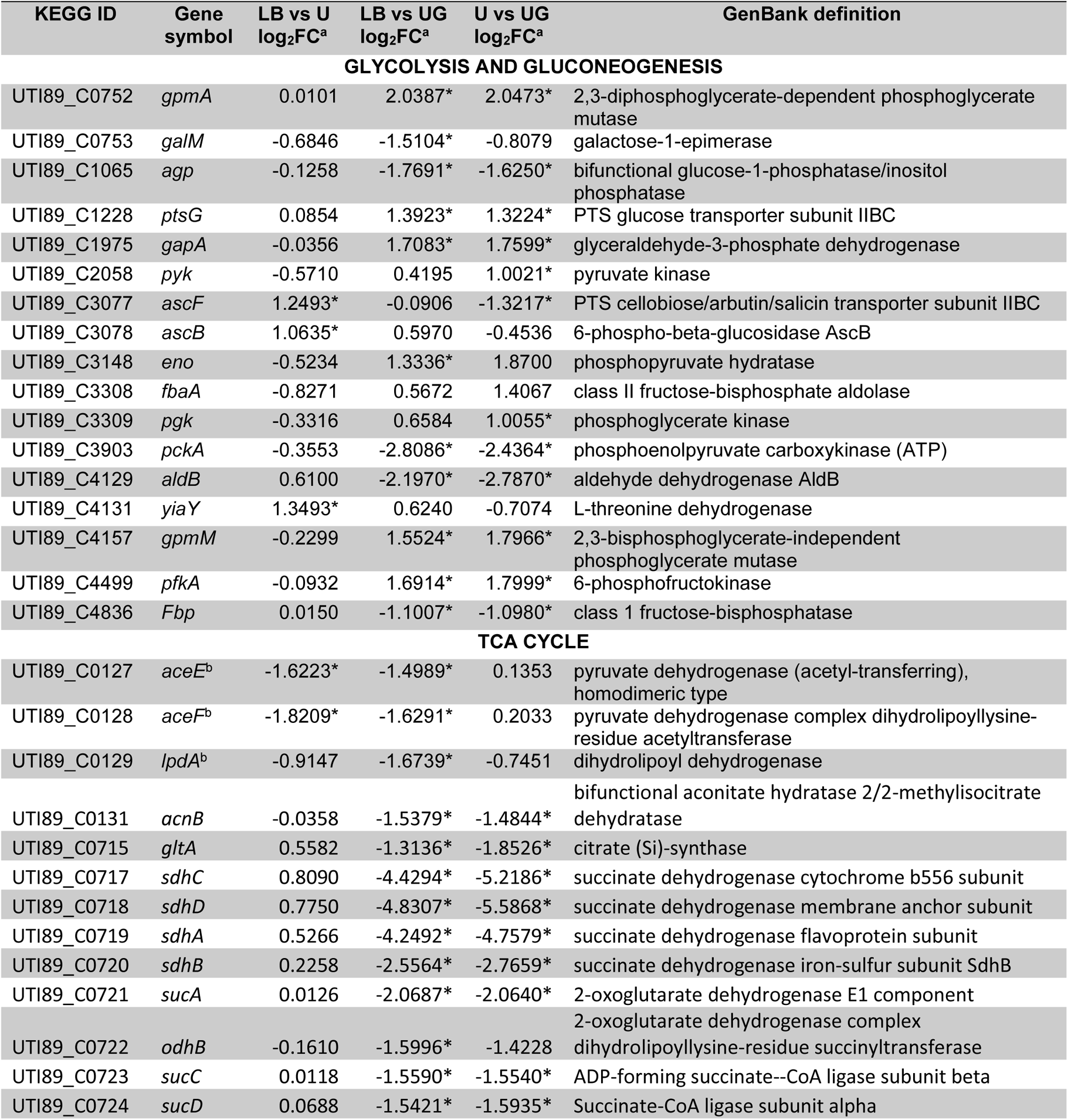
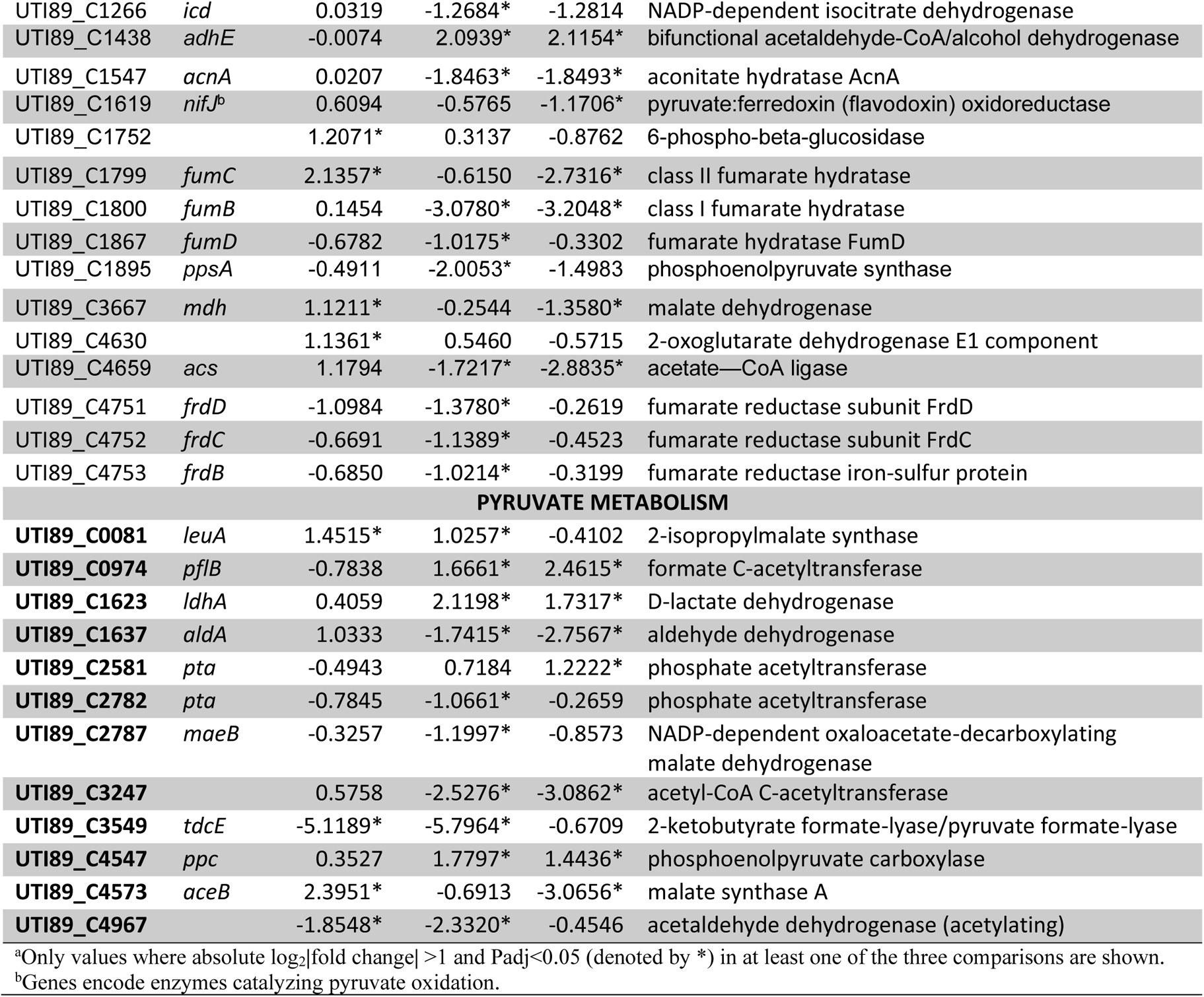
RNA-Seq analysis of expression of genes involved in glucose metabolism

| KEGG ID | Gene symbol | LB vs U<br>log<sub>2</sub>FC<sup>a</sup> | LB vs UG<br>log<sub>2</sub>FC<sup>a</sup> | U vs UG<br>log<sub>2</sub>FC<sup>a</sup> | GenBank definition |
| --- | --- | --- | --- | --- | --- |
| **GLYCOLYSIS AND GLUCONEOGENESIS** |  |  |  |  |  |
| UTI89_C0752 | *gpmA* | 0.0101 | 2.0387* | 2.0473* | 2,3-diphosphoglycerate-dependent phosphoglycerate mutase |
| UTI89_C0753 | *galM* | -0.6846 | -1.5104* | -0.8079 | galactose-1-epimerase |
| UTI89_C1065 | *agp* | -0.1258 | -1.7691* | -1.6250* | bifunctional glucose-1-phosphatase/inositol phosphatase |
| UTI89_C1228 | *ptsG* | 0.0854 | 1.3923* | 1.3224* | PTS glucose transporter subunit IIBC |
| UTI89_C1975 | *gapA* | -0.0356 | 1.7083* | 1.7599* | glyceraldehyde-3-phosphate dehydrogenase |
| UTI89_C2058 | *pyk* | -0.5710 | 0.4195 | 1.0021* | pyruvate kinase |
| UTI89_C3077 | *ascF* | 1.2493* | -0.0906 | -1.3217* | PTS cellobiose/arbutin/salicin transporter subunit IIBC |
| UTI89_C3078 | *ascB* | 1.0635* | 0.5970 | -0.4536 | 6-phospho-beta-glucosidase AscB |
| UTI89_C3148 | *eno* | -0.5234 | 1.3336* | 1.8700 | phosphopyruvate hydratase |
| UTI89_C3308 | *fbaA* | -0.8271 | 0.5672 | 1.4067 | class II fructose-bisphosphate aldolase |
| UTI89_C3309 | *pgk* | -0.3316 | 0.6584 | 1.0055* | phosphoglycerate kinase |
| UTI89_C3903 | *pckA* | -0.3553 | -2.8086* | -2.4364* | phosphoenolpyruvate carboxykinase (ATP) |
| UTI89_C4129 | *aldB* | 0.6100 | -2.1970* | -2.7870* | aldehyde dehydrogenase AldB |
| UTI89_C4131 | *yiaY* | 1.3493* | 0.6240 | -0.7074 | L-threonine dehydrogenase |
| UTI89_C4157 | *gpmM* | -0.2299 | 1.5524* | 1.7966* | 2,3-bisphosphoglycerate-independent phosphoglycerate mutase |
| UTI89_C4499 | *pfkA* | -0.0932 | 1.6914* | 1.7999* | 6-phosphofructokinase |
| UTI89_C4836 | *Fbp* | 0.0150 | -1.1007* | -1.0980* | class 1 fructose-bisphosphatase |
| **TCA CYCLE** |  |  |  |  |  |
| UTI89_C0127 | *aceE<sup>b</sup>* | -1.6223* | -1.4989* | 0.1353 | pyruvate dehydrogenase (acetyl-transferring), homodimeric type |
| UTI89_C0128 | *aceF<sup>b</sup>* | -1.8209* | -1.6291* | 0.2033 | pyruvate dehydrogenase complex dihydrolipoyllysine-residue acetyltransferase |
| UTI89_C0129 | *lpdA<sup>b</sup>* | -0.9147 | -1.6739* | -0.7451 | dihydrolipoyl dehydrogenase |
| UTI89_C0131 | *acnB* | -0.0358 | -1.5379* | -1.4844* | bifunctional aconitate hydratase 2/2-methylisocitrate dehydratase |
| UTI89_C0715 | *gltA* | 0.5582 | -1.3136* | -1.8526* | citrate (Si)-synthase |
| UTI89_C0717 | *sdhC* | 0.8090 | -4.4294* | -5.2186* | succinate dehydrogenase cytochrome b556 subunit |
| UTI89_C0718 | *sdhD* | 0.7750 | -4.8307* | -5.5868* | succinate dehydrogenase membrane anchor subunit |
| UTI89_C0719 | *sdhA* | 0.5266 | -4.2492* | -4.7579* | succinate dehydrogenase flavoprotein subunit |
| UTI89_C0720 | *sdhB* | 0.2258 | -2.5564* | -2.7659* | succinate dehydrogenase iron-sulfur subunit SdhB |
| UTI89_C0721 | *sucA* | 0.0126 | -2.0687* | -2.0640* | 2-oxoglutarate dehydrogenase E1 component |
| UTI89_C0722 | *odhB* | -0.1610 | -1.5996* | -1.4228 | 2-oxoglutarate dehydrogenase complex dihydrolipoyllysine-residue succinyltransferase |
| UTI89_C0723 | *sucC* | 0.0118 | -1.5590* | -1.5540* | ADP-forming succinate--CoA ligase subunit beta |
| UTI89_C0724 | *sucD* | 0.0688 | -1.5421* | -1.5935* | Succinate-CoA ligase subunit alpha |
| UTI89_C1266 | *icd* | 0.0319 | -1.2684* | -1.2814 | NADP-dependent isocitrate dehydrogenase |
| UTI89_C1438 | *adhE* | -0.0074 | 2.0939* | 2.1154* | bifunctional acetaldehyde-CoA/alcohol dehydrogenase |
| UTI89_C1547 | *acnA* | 0.0207 | -1.8463* | -1.8493* | aconitate hydratase AcnA |
| UTI89_C1619 | *nifJ*<sup>b</sup> | 0.6094 | -0.5765 | -1.1706* | pyruvate:ferredoxin (flavodoxin) oxidoreductase |
| UTI89_C1752 |  | 1.2071* | 0.3137 | -0.8762 | 6-phospho-beta-glucosidase |
| UTI89_C1799 | *fumC* | 2.1357* | -0.6150 | -2.7316* | class II fumarate hydratase |
| UTI89_C1800 | *fumB* | 0.1454 | -3.0780* | -3.2048* | class I fumarate hydratase |
| UTI89_C1867 | *fumD* | -0.6782 | -1.0175* | -0.3302 | fumarate hydratase FumD |
| UTI89_C1895 | *ppsA* | -0.4911 | -2.0053* | -1.4983 | phosphoenolpyruvate synthase |
| UTI89_C3667 | *mdh* | 1.1211* | -0.2544 | -1.3580* | malate dehydrogenase |
| UTI89_C4630 |  | 1.1361* | 0.5460 | -0.5715 | 2-oxoglutarate dehydrogenase E1 component |
| UTI89_C4659 | *acs* | 1.1794 | -1.7217* | -2.8835* | acetate—CoA ligase |
| UTI89_C4751 | *frdD* | -1.0984 | -1.3780* | -0.2619 | fumarate reductase subunit FrdD |
| UTI89_C4752 | *frdC* | -0.6691 | -1.1389* | -0.4523 | fumarate reductase subunit FrdC |
| UTI89_C4753 | *frdB* | -0.6850 | -1.0214* | -0.3199 | fumarate reductase iron-sulfur protein |
| **PYRUVATE METABOLISM** |  |  |  |  |  |
| UTI89_C0081 | *leuA* | 1.4515* | 1.0257* | -0.4102 | 2-isopropylmalate synthase |
| UTI89_C0974 | *pflB* | -0.7838 | 1.6661* | 2.4615* | formate C-acetyltransferase |
| UTI89_C1623 | *ldhA* | 0.4059 | 2.1198* | 1.7317* | D-lactate dehydrogenase |
| UTI89_C1637 | *aldA* | 1.0333 | -1.7415* | -2.7567* | aldehyde dehydrogenase |
| UTI89_C2581 | *pta* | -0.4943 | 0.7184 | 1.2222* | phosphate acetyltransferase |
| UTI89_C2782 | *pta* | -0.7845 | -1.0661* | -0.2659 | phosphate acetyltransferase |
| UTI89_C2787 | *maeB* | -0.3257 | -1.1997* | -0.8573 | NADP-dependent oxaloacetate-decarboxylating malate dehydrogenase |
| UTI89_C3247 |  | 0.5758 | -2.5276* | -3.0862* | acetyl-CoA C-acetyltransferase |
| UTI89_C3549 | *tdcE* | -5.1189* | -5.7964* | -0.6709 | 2-ketobutyrate formate-lyase/pyruvate formate-lyase |
| UTI89_C4547 | *ppc* | 0.3527 | 1.7797* | 1.4436* | phosphoenolpyruvate carboxylase |
| UTI89_C4573 | *aceB* | 2.3951* | -0.6913 | -3.0656* | malate synthase A |
| UTI89_C4967 |  | -1.8548* | -2.3320* | -0.4546 | acetaldehyde dehydrogenase (acetylating) |

<sup>a</sup>Only values where absolute log<sub>2</sub>|fold change| >1 and Padj < 0.05 (denoted by \*) in at least one of the three comparisons are shown.
<sup>b</sup>Genes encode enzymes catalyzing pyruvate oxidation.

**Table 3:**
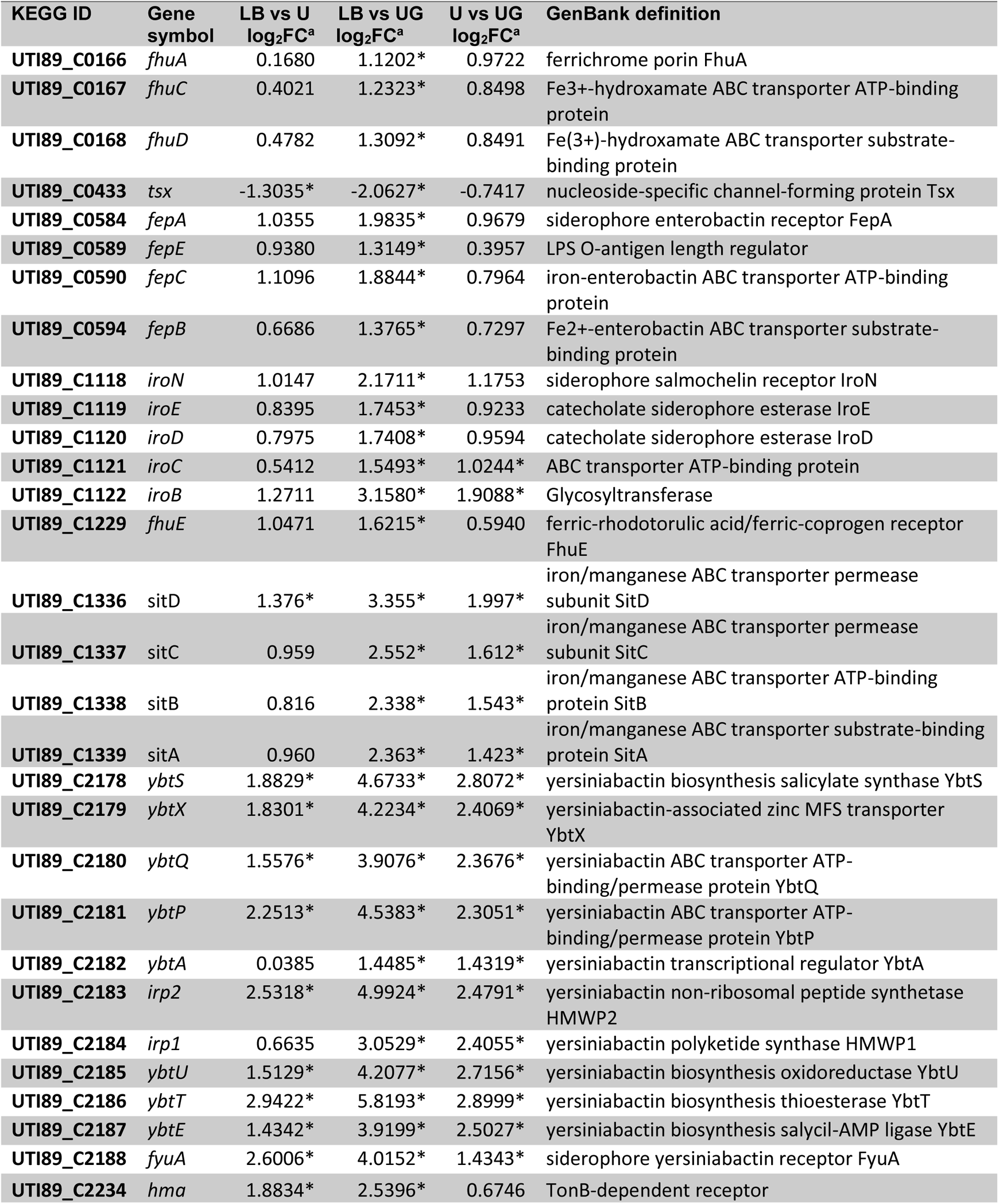
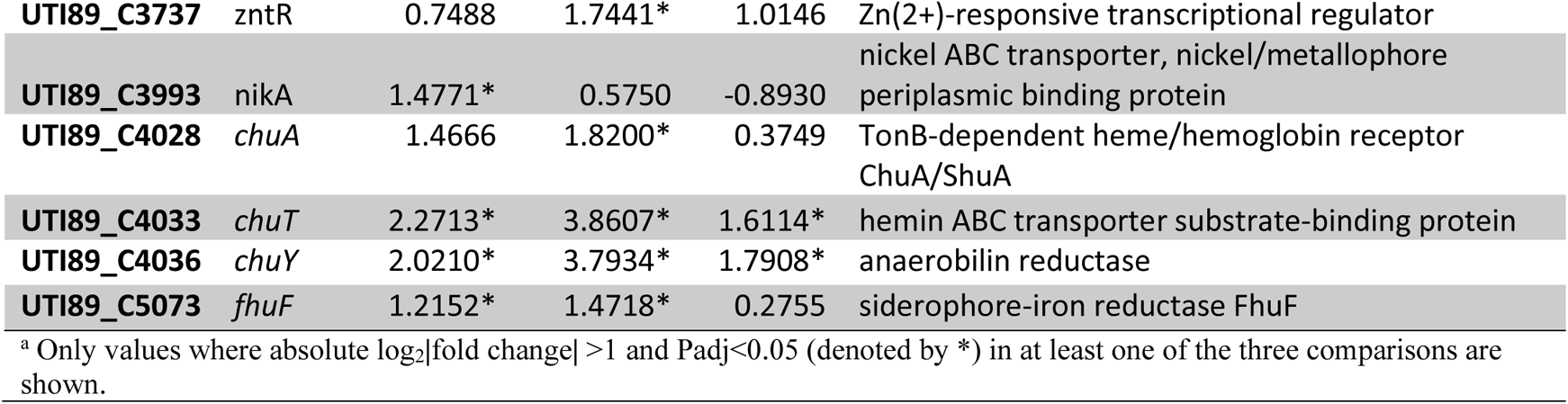
RNA-Seq analysis of expression of genes involved in metal transport

**Table 4:** RNA-Seq results for virulence genes^.

| KEGG ID | Gene symbol | LB vs U<br>log<sub>2</sub>FC<sup>a</sup> | LB vs UG<br>log<sub>2</sub>FC<sup>a</sup> | U vs UG<br>log<sub>2</sub>FC<sup>a</sup> | GenBank Definition |
| --- | --- | --- | --- | --- | --- |
| UTI89_C0149 |  | 1.0177 | -0.0745 | -1.0715* | fimbrial-like adhesin |
| UTI89_C0152 |  | 0.7837 | -0.9048 | -1.6806* | fimbrial protein |
| UTI89_C0154 |  | 0.9027 | 1.1416* | 0.2621 | fimbrial chaperone |
| UTI89_C0155 |  | 0.3290 | 1.3269* | 1.0121 | fimbrial protein |
| UTI89_C1110 | *sfaD* | -0.2407 | -1.2714* | -1.0148 | type 1 fimbrial protein |
| UTI89_C1111 |  | -0.0963 | -1.4300* | -1.3136 | S/F1C fimbrial biogenesis chaperone SfaE/FocC |
| UTI89_C1114 |  | -0.3881 | -1.0051* | -0.5956 | S-fimbrial adhesin minor pilin SfaS |
| UTI89_C1139 | *ag43* | -1.5590* | -1.1735* | 0.4048 | autotransporter adhesin Ag43 |
| UTI89_C1159 | *csgF* | 1.1020 | -1.2543* | -2.3179* | curli production assembly/transport protein CsgF |
| UTI89_C1160 | *csgE* | 0.9886 | -1.2301 | -2.2080* | curli production assembly/transport protein CsgE |
| UTI89_C1161 | *csgD* | 1.1839 | -0.8703 | -2.0304* | transcriptional regulator CsgD |
| UTI89_C1719 |  | -1.9731* | -2.6289* | -0.6357 | fimbrial protein |
| UTI89_C2094 | *flhC* | 0.6029 | -1.1429 | -1.7342* | flagellar transcriptional regulator FlhC |
| UTI89_C2095 | *flhD* | 0.7667 | -1.4090* | -2.1631* | flagellar transcriptional regulator FlhD |
| UTI89_C2151 | *rcsA* | -0.1543 | -1.1939* | -1.0163* | transcriptional regulator RcsA |
| UTI89_C3049 | *luxS* | -0.3816 | 0.6139 | 1.0096* | S-ribosylhomocysteine lyase |
| UTI89_C3180 | *gcvA* | 1.2075* | 1.3061* | 0.1140 | glycine cleavage system transcriptional regulator GcvA |
| UTI89_C3362 | *kpsF* | -1.0395 | -2.1530* | -1.0959 | KpsF/GutQ family sugar-phosphate isomerase |
| UTI89_C3363 | *kpsE* | -0.6047 | -1.1880* | -0.5649 | capsule polysaccharide export inner-membrane protein KpsE |
| UTI89_C3364 | *kpsD* | -0.7698 | -1.0469* | -0.2603 | polysialic acid transporter KpsD |
| UTI89_C3367 |  | -0.7502 | -1.3350* | -0.5646 | capsular biosynthesis protein |
| UTI89_C3935 |  | -0.0955 | -1.1975* | -1.0802* | fimbria/pilus periplasmic chaperone |
| UTI89_C4639 | *kguS* | 0.4967 | -1.0320* | -1.4958* | sensor histidine kinase |
| UTI89_C4907 |  | -1.1595* | -0.5895 | 0.5885 | fimbrial protein |
| UTI89_C4926 | *hlyA* | -1.4809* | -1.8559* | -0.3579 | RTX toxin hemolysin HlyA |
| UTI89_C4927 | *hlyC* | -1.6279* | -2.0831* | -0.4356 | alpha-hemolysin-activating lysine-acyltransferase HlyC |
| UTI89_C5009 | *fimB* | -1.9050* | -2.6000* | -0.6761 | type 1 fimbria switch DNA invertase FimB |
| UTI89_C5010 | *fimE* | 1.9997* | 1.7552* | -0.2264 | type 1 fimbria switch DNA invertase FimE |
| **D-SERINE AND L-SERINE METABOLISM<sup>^</sup>** |  |  |  |  |  |
| UTI89_C2010 | *sdaA* | -0.1862 | -1.4675* | -1.4675* | L-serine ammonia-lyase |
| UTI89_C3168 | *sdaB* | -0.6305 | -2.2129* | -2.2129* | L-serine ammonia-lyase II |
| UTI89_C3167 | *sdaC* | -0.2304 | -2.8554* | -2.8554* | HAAAP family serine/threonine permease SdaC |
| UTI89_C4955 | *dsdA* | 1.1998* | 0.3089 | -0.8768 | D-serine ammonia-lyase |
| **UTI89_C4956** | dsdX | 1.2850* | 0.5240 | -0.7400 | D-serine transporter DsdX |
| **UTI89_C4957** | dsdC | 1.3554* | 0.4634 | -0.8736 | DNA-binding transcriptional regulator DsdC |

^ includes genes from KEGG pathway “biofilm formation” (*eci02026*) and genes involved in metabolism and transport of D-serine (*dsdA/X/C*) and L-serine (*sdaA/B/C*), shown to play a role in the uropathogenesis. <sup>a</sup> Only values where absolute log<sub>2</sub>[fold change] >1 and Padj<0.05 (denoted by \*) in at least one of the three comparisons are shown.

**Table 5:** RNA-Seq results for genes from category ‘human diseases’ in KEGG database

| KEGG ID | Gene symbol | LB vs U<br>log<sub>2</sub>FC<sup>a</sup> | LB vs UG<br>log<sub>2</sub>FC<sup>a</sup> | U vs UG<br>log<sub>2</sub>FC<sup>a</sup> | GenBank Definition |
| --- | --- | --- | --- | --- | --- |
| **Beta-lactam resistance** |  |  |  |  |  |
| UTI89_C0275 |  | 0.1184 | -1.0975* | -1.1988* | RNA ligase RtcB family protein |
| UTI89_C0457 | *ampG* | -0.2442 | -1.3200* | -1.0604 | muropeptide MFS transporter AmpG |
| UTI89_C1001 | *ompF* | 0.1189 | -2.1144* | -2.2156* | porin OmpF |
| UTI89_C1601 | *mppA* | 0.1456 | -1.3388* | -1.4675* | murein tripeptide ABC transporter substrate-binding protein MppA |
| UTI89_C2497 | *ompC* | -1.2018* | -0.6919 | 0.5258 | porin OmpC |
| **Vancomycin resistance** |  |  |  |  |  |
| UTI89_C0095 | *murF* | -0.7417 | 0.3595 | 1.1134* | UDP-N-acetylmuramoyl-tripeptide--D-alanyl-D-alanine ligase |
| UTI89_C0096 | *mraY* | -0.9774 | 0.1922 | 1.1833* | phospho-N-acetylmuramoyl-pentapeptide-transferase |
| UTI89_C0099 | *murG* | -1.2970* | -0.0013 | 1.3099* | undecaprenyldiphospho-muramoylpentapeptide beta-N-acetylglucosaminyltransferase |
| UTI89_C0100 | *murC* | -1.2638* | -0.0551 | 1.2231 | UDP-N-acetylmuramate--L-alanine ligase |
| UTI89_C1376 | *dadX* | 0.0759 | -1.5158* | -1.5720* | catabolic alanine racemase DadX |
| **CAMP resistance** |  |  |  |  |  |
| UTI89_C0626 | *pagP* | -0.8325 | 0.3986 | 1.2467* | lipid IV(A) palmitoyltransferase PagP |
| UTI89_C1750 | *marA* | 3.2992* | 1.7309* | -1.5534* | MDR efflux pump AcrAB transcriptional activator MarA |
| UTI89_C2535 | *arnB* | -0.6266* | -1.2033* | -0.5596 | UDP-4-amino-4-deoxy-L-arabinose aminotransferase |
| UTI89_C2536 | *arnC* | -0.7728 | -1.2099* | -0.4226 | undecaprenyl-phosphate 4-deoxy-4-formamido-L-arabinose transferase |
| UTI89_C4708 | *eptA* | -2.2420* | -2.2556* | 0.0061 | phosphoethanolamine transferase EptA |

<sup>a</sup> Only values where absolute log<sub>2</sub>[fold change] >1 and Padj<0.05 (denoted by \*) in at least one of the three comparisons are shown.

As healthy urine is devoid of free carbohydrates, the principal source of carbon in the urinary tract is small amounts of amino acids and peptides (34). During colonization of a healthy urinary tract (without glycosuria), following their trans-membrane transport, amino acids are catabolized by UPEC via TCA cycle and gluconeogenesis to synthesize glucose. In presence of glycosuria, however, central carbon metabolism is expected to be routed through glycolysis. We examined RNASeq data for peptide and amino acid import (Table 1), glucose metabolism (glycolysis, gluconeogenesis, TCA cycle, and pyruvate metabolism genes, Table 2).

Compared to UTI89-LB, both UTI89-U and UTI89-UG showed upregulation of genes encoding transporters of serine (*dsdX*, UTI89_C4956; *cycA*, UTI89_C4817; *sstT*, UTI89_C3527), arginine (*artJ/M/Q/J/P*, UTI89_C0863—67), histidine (*hisQ/J*, UTI89_C2592/93), branched chain amino acids (*livF/G/M/H/K/J*, UTI89_C3961—65, C3969) and dipeptides (*dppB*/*A*, UTI89_C4081/82). In contrast glycine/proline ABC transporter (*proV/W/X*, UTI89_C3037—39) genes were significantly downregulated in UTI89-U and UTI89-UG compared to UTI89-LB. Differential expression of genes involved in glucose metabolism (*eci00010*-glycolysis/gluconeogenesis, *eci00020*-TCA cycle, *eci00620*-pyruvate metabolism) is presented in table 2. In UTI89-UG, we observed significant upregulation of glycolysis genes *gpmA* (UTI89_C0752), *gapA* (UTI89_C1975), *pfkA* (UTI89_C2058), *pgk* (UTI89_C3309), *gpmM* (UTI89_C4157), and *pyk* (UTI89_C4499). UTI89-UG also showed significant downregulation of gluconeogenesis genes *pckA* (UTI89_C3903, pyruvate carboxykinase) and *fbp* (UTI89_4836, fructose 1,6 bisphosphatase) encoding enzymes that catalyze bypass of irreversible steps of glycolysis and of *fumB* (UTI89_C1800, fumarate hydratase) catalyzing conversion of malate to fumarate in reverse TCA cycle. Overall, these results indicate that exposure to glycosuria induces metabolic transition of UPEC towards glycolysis. In contrast, UTI89-U exhibited significant upregulation of *fumC* (UTI89_1799, fumarate hydratase catalyzes fumarate oxidation to malate) and *mdh* (UTI89_ 3667, malate dehydrogenase catalyzes malate to oxaloacetate), while expression of gluconeogenesis genes *pckA*, *fbp*, or *fumB* was not significantly altered in UTI89-U. Serine, present in high amount in urine, is preferentially catabolized by UPEC into acetate (34, 36). For the genes encoding enzymes involved in serine to acetate catabolism, we observed significant downregulation of phosphate acetyl transferase (*pta*, UTI89_C2782) in UTI89-UG while acetate kinase (*ackA*, UTI89_C2579) was unchanged in either UTI89-U or UTI89-UG. UTI89-UG also showed significant downregulation of acetate-coA ligase (*acs*, UTI89_C4659) involved in acetate utilization (37).

Next, we examined RNASeq data for genes encoding metal ion transporters (iron, manganese, nickel, cobalt, copper, zinc), which constitute important determinants of urinary fitness and virulence of UPEC (Table 4). Exposure to glycosuria (UTI89-UG) showed upregulation of iron uptake systems importing siderophores such as yersiniabactin (*ybt*/*fyuA*, UTI89_C2178— 87/C2188), enterobactin (*fepA*/*E*/*C*/*B* (UTI89_C0584/89/90/94), and salmochelin (*iroN*/*E*/*D*/*C*/*B*, UTI89_C1118—22), ferric hydroxamate transporter (*fhuA*/*C*/*D*/*E*, UTI89_C0166/67/68/C1229), free ferrous/manganese ion importer *sitD*/*C*/*B*/*A* (UTI89_C1336—39), *chuA*/*T*/*Y* (UTI89_C4028/4033/4036) importer of iron bound to host protein heme. UTI89-UG versus UTI89-LB also showed upregulation of *zntR* (UTI89_C3737) zinc transporter. In UTI89-U versus UTI89-LB comparison we observed significant upregulation of encoding *ybt* yersiniabactin iron acquisition system, and *chu* heme uptake system, and nickel transporter *nikA* (UTI89_C3993). We did not observe changes in the expression of copper export genes (*copA*/*D* and *cusS*/*R*/*C*/*B*/*A*) in UTI89-U or UTI89-UG. Biofilm formation in *E. coli* is primarily regulated via secondary messenger cAMP (3’,5’ cyclic adenosine monophosphate)—CRP (cAMP receptor protein) complex. cAMP-CRP complex induces biosynthesis of surface adhesive type 1 pili, curli fibers, flagella, and K1 capsule and suppresses sigma factor *RpoS*, a known repressor of initiation of biofilms (38, 39). Glucose-mediated catabolite repression suppresses *E. coli* biofilms by downregulation of *cyaA* encoding enzyme adenylyl cyclase which catalyzes cAMP production (38). Indeed, in UTI89-UG we observed significant downregulation of positive regulators of type 1 pilus, curli fiber, and flagellar biosynthesis, *fimB* (UTI89_C5009), *csgD* (UTI89_C1161), and *flhC*/*D* (UTI89_C2094/C2094), respectively (Table 4). We did not observe significant changes in the expression of *csrA* (UTI89_C3057 encodes an RNA binding protein that disseminates biofilms)*, cyaA* (UTI89_4365)*, or crp* (UTI89_C3860) in any of the three transcriptome comparisons. UTI89-UG also showed downregulation of *kpsF/E/D* (UTI89_C3362/63/64) encoding K1 capsule. UTI89-U and UTI89-UG also showed significant downregulation of *ag43* (UTI89_C1139) which induces autoaggregation, biofilm formation, and long-term persistence of UPEC in a mouse model of ascending UTI (40). UTI89-UG showed significant reduction in the expression in hemolysin genes *hlyA/C* (UTI89_C4926/27) shown to activate bladder cell exfoliation via caspase-mediated pro-inflammatory cell death (41) and in the expression of two component system genes involved in *α*-ketoglutarate utilization (*kguS*, UTI89_C4639) critical for nutritional fitness of UPEC in the urinary tract (42).

Table 5 shows differential expression analysis for genes categorized under ‘human diseases’ for KEGG pathways beta-lactam resistance (*eci01501*), Vancomycin resistance (eci1502), cationic antimicrobial peptide (CAMP) resistance (eci1503). Compared to UTI89-U, UTI89-UG showed significantly increased expression of vancomycin resistance genes *murF* (UTI89_C0095), *mraY* (UTI89_C0096), *murG* (UTI89_C0099), and *murC* (UTI89_C0101) and CAMP resistance gene *pagP* (UTI89_C0626). Mur enzymes (MurABCDEF) catalyze synthesis of UDP *N*-acetyl muramyl pentapeptide from UDP *N*-acetyl glucosamine, a major component of UPEC cell wall peptidoglycan, which determines bacterial shape and mediates protection.

Of these, *murCDEF* were significantly upregulated in UTI89-UG compared to UTI89-U. In contrast, β-lactam resistance genes *ampG* (UTI89_C0275, UTI89_C0457), *ompF* (UTI89_C1001), and *mppA* (UTI89_C1601) were significantly downregulated in UTI89-UG compared to UTI-U.

### qRTPCR analysis of UTI89 gene expression

Using qRTPCR, we examined mRNA transcript levels in UTI89-LB, UTI89-U, UTI89-UG, and UTI89 UGal for a panel of genes encoding *hlyA* (UTI89_C4926) and iron acquisition proteins, *chuT* (UTI89_C4033), *fur* (UTI89_C0687), *fyuA* (UTI89_C2188), *hma* (UTI89_C2234), and *irp2* (UTI89_C2183). Fold difference in specific mRNA transcript levels (relative to housekeeping gene transcripts, relative quantification or RQ) over RQ for UTI89-LB control are presented in Fig 2. As mentioned earlier, comparisons UTI-LB versus UTI89-U and UTI89-LB versus UTI89-UG showed significant correlation with RNASeq results across the genes (supplementary Fig S4). Importantly, gene expression profile for UTI89 exposed for 2h to urine supplemented with galactose (Fig 2B) was similar to that observed in UTI89-UG (UTI89-UGal) suggesting that osmolarity change may be responsible for differential gene expression at least for the genes in our panel.

### Glycosuria induces UPEC biofilm formation

Biofilms are multi-bacterial communities organized in a three-dimensional space in a matrix made of proteins, polysaccharides, and DNA secreted by biofilm-bound bacteria (43). UPEC produces biofilms on the surface and inside bladder epithelial cells as well as on urinary catheters (44). To determine the effects of glycosuria on biofilm formation, type 1 piliated UPEC were washed in PBS and resuspended in LB, U, or UG for biofilm formation in a 96-well, flat-bottom, polystyrene plate. After overnight incubation at 37°C and 150rpm, biofilms were examined by crystal violet staining as well as by enumerating planktonic (supernatant) and biofilm-bound CFUs. We observed that glycosuria (UG) significantly augments biofilm formation by both UTI89 and CFT073, as adjudged by more intense violet staining of biofilm biomass in UTI89-UG (Fig 4 A) and CFT073-UG (Fig 4 C) compared to their counterparts cultivated in either LB or U. UTI89-UG also showed significantly higher proportion of biofilm-bound CFUs in comparison to LB-control and UTI89-U (Fig 4B), while increased proportion of CFT073-UG CFUs was not statistically significant in comparison to either LB-control or CFT073-U. Urine alone did not induce biofilm formation in either UTI89 or CFT073 compared to LB controls. These results suggest that glycosuria augments UPEC biofilms, which was surprising as the previous research has shown that glucose-mediated catabolite repression prevent biofilm formation via cAMP-CRP (38, 39). Iron availability is linked with biofilm formation via the activation of *ybt-fyuA* yersiniabactin iron uptake system as UPEC mutant ablated for fyuA activity shows reduced biofilm formation (23). We also observed that exposure to glycosuria (UTI89-UG) induces expression of *ybt-fyuA* yersiniabactin gene cluster (Table 4). Hence we examined the effects of iron chelation on glycosuria-induced UPEC biofilms by supplementing LB, U, and UG with a membrane permeable iron chelator, DIP. Addition of DIP did not reduce augmentation of either UTI89 or CFT073 biofilms in the presence of UG (data not shown). Future experiments comparing transcriptomic profiles of UPEC collected at different timepoints (6, 12, and 24h) during biofilm formation in LB, U, or UG will reveal potential effectors involved in the formation of UPEC biofilms.

**Figure 4:**
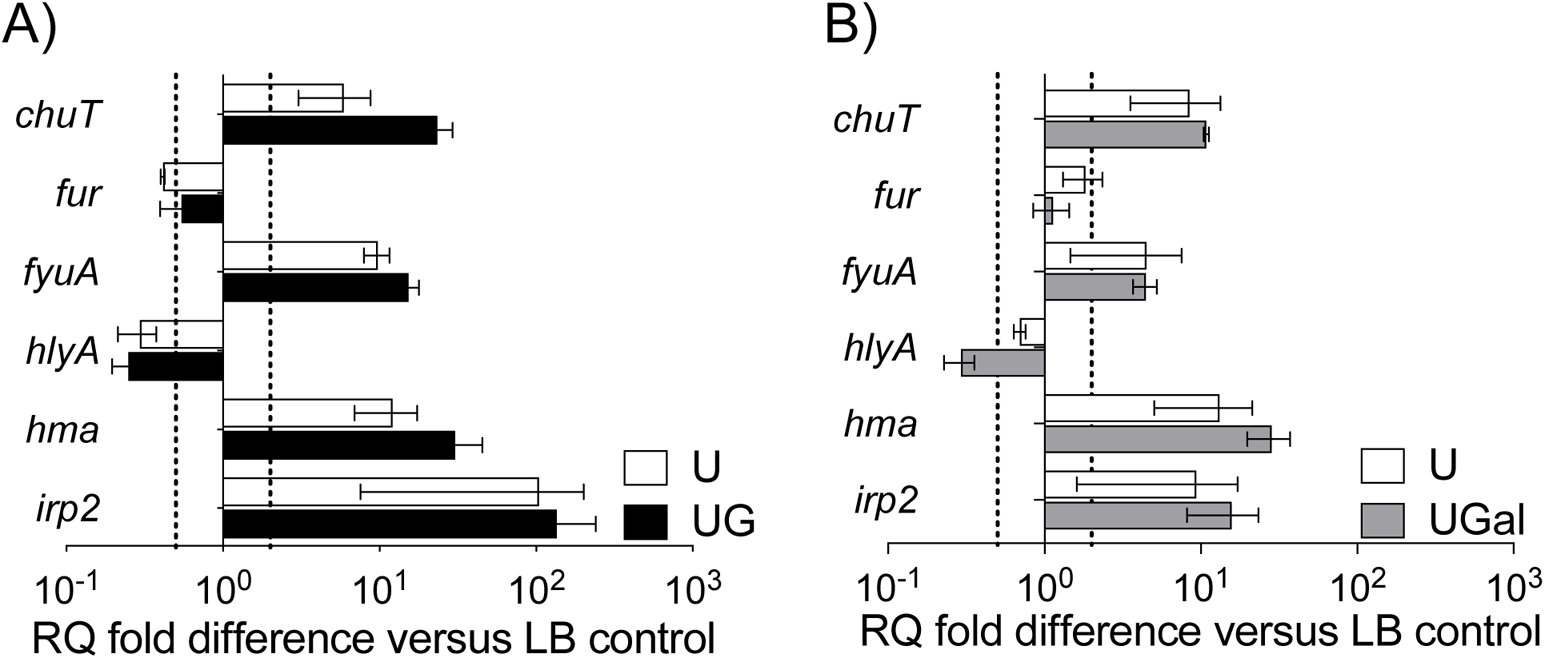
qRTPCR results for UPEC strains exposed to urine plain or supplemented with glucose or galactose. Using quantitative real time PCR (qRTPCR) with normalization to 16SrRNA to determine mRNA transcript levels for specific virulence and associated genes (indicated on *x*-axis) in UTI89 following 2h-long exposure to U or UG (A) or to U or UGal (B). RQ values were calculated by comparative threshold cycle (*ΔΔ*C_T_) algorithm. RQ fold differences over transcript levels from LB-control are presented as average of at least two biological replicates (each with three technical repeats) ± standard deviation. Dotted lines indicate 2-fold up—or down—regulation.

Next, we examined whether increase in urine osmolality due to the presence of glucose plays a role in this phenotype by measuring biofilm formation in urine supplemented with 600mg/dl of galactose (UGal). The presence of equal amount (600mg/dl) of either glucose or galactose alters urine osmolality by a similar extent. Compared to LB-control, both UTI89-UGal and CFT073-Ugal showed significantly higher biofilm biomass by crystal violet staining (Fig 4A, C). CFU enumeration showed increased proportion of biofilm-bound CFUs for UTI89-UGal compared to UTI89-LB or UTI89-U; the proportion of biofilm CFUs in UTI89-UGal was significantly lower than that in UTI89-UG (Fig 4B). Interestingly, biofilm-bound CFUs in CFT073-UGal were similar to those in CFT073 in LB or U; while proportion of biofilm CFUs in CFT073-UGal was lower than that in CFT073-UG although this difference was not statistically significant (Fig 4D). Overall, these results suggest that urine osmolarity plays a role in glycosuria-mediated augmentation of UPEC biofilms.

### Glycosuria affects UPEC hemagglutination of guinea pig RBCs

UPEC adherence to bladder epithelium mediated by type 1 pili is the first step in colonization of the urinary tract during ascending UTI. Guinea pig hemagglutination (HA) assays for the assembly of functional type 1 pili on the bacterial surface, where guinea pig RBCs are mixed with two-fold serial dilution of UPEC. HA titer is reported as the lowest bacterial dilution at which UPEC is able to agglutinate guinea pig RBCs. We observed that compared to LB-control (set to 100% HA titer), HA titers of UTI89-U and CFT073-U were ~2-fold decreased, while HA titers for UTI89-UG and CFT073-UG were reduced by ~10-fold and 5-fold respectively. HA results correlate with previous observations that glucose suppresses type 1 piliation in UPEC via cAMP-CRP complex (45)

### Glycosuria induces UPEC virulence in a mouse model of ascending UTI

To examine the effects of glycosuria on UPEC virulence, we experimentally induced ascending UTI in 8 weeks old male and female C3H mice by inoculating via transurethral catheterization UTI89 or CFT073 pre-exposed (2h, 37°C, static) to either LB, U, or UG. Bladder, kidneys, and spleen homogenates were dilution plated at 24hpi to enumerate bacterial organ burden. No bacteria were recovered from the spleen, indicating lack of dissemination via blood. The median bladder CFU burden in mice infected with UTI89-UG was ~4-fold higher (*P*= 0.27, Mann Whitney U test) compared to those infected with UTI89-LB and 200-fold higher (*P*=0.003) compared to mice infected with UTI89-U (Fig 5A). The median kidney CFUs of mice infected with UTI89-UG showed no change compared to those infected with UTI89-LB and ~4-fold increase (*P*=0.019) compared to mice infected with UTI89-U (Fig 5A). The median bladder CFU burden in mice infected with CFT073-UG showed ~3-fold increase compared to mice infected with either CFT073-LB (*P*= 0.8, U test) or CFT073-U (*P*=0.6; Fig 5A). The median kidney CFUs in mice infected with CFT073-UG showed 7.5-fold decrease compared to those infected with CFT073-LB (*P*= 0.07, U test) and 2.5-fold decrease compared to mice infected with CFT073-U (*P*=0.03; Fig 5B). Interestingly, compared to their UTI89-LB counterparts, mice infected with UTI89-U showed 55-fold (*P*=0.054) reduction in median bladder CFUs and 4-fold (*P*=0.01) reduction in median kidney CFUs. CFT073-U infected mice also showed 10-fold reduction (*P*=0.6) in median bladder CFUs compared to CFT073-LB infected counterparts.

**Figure 5:**
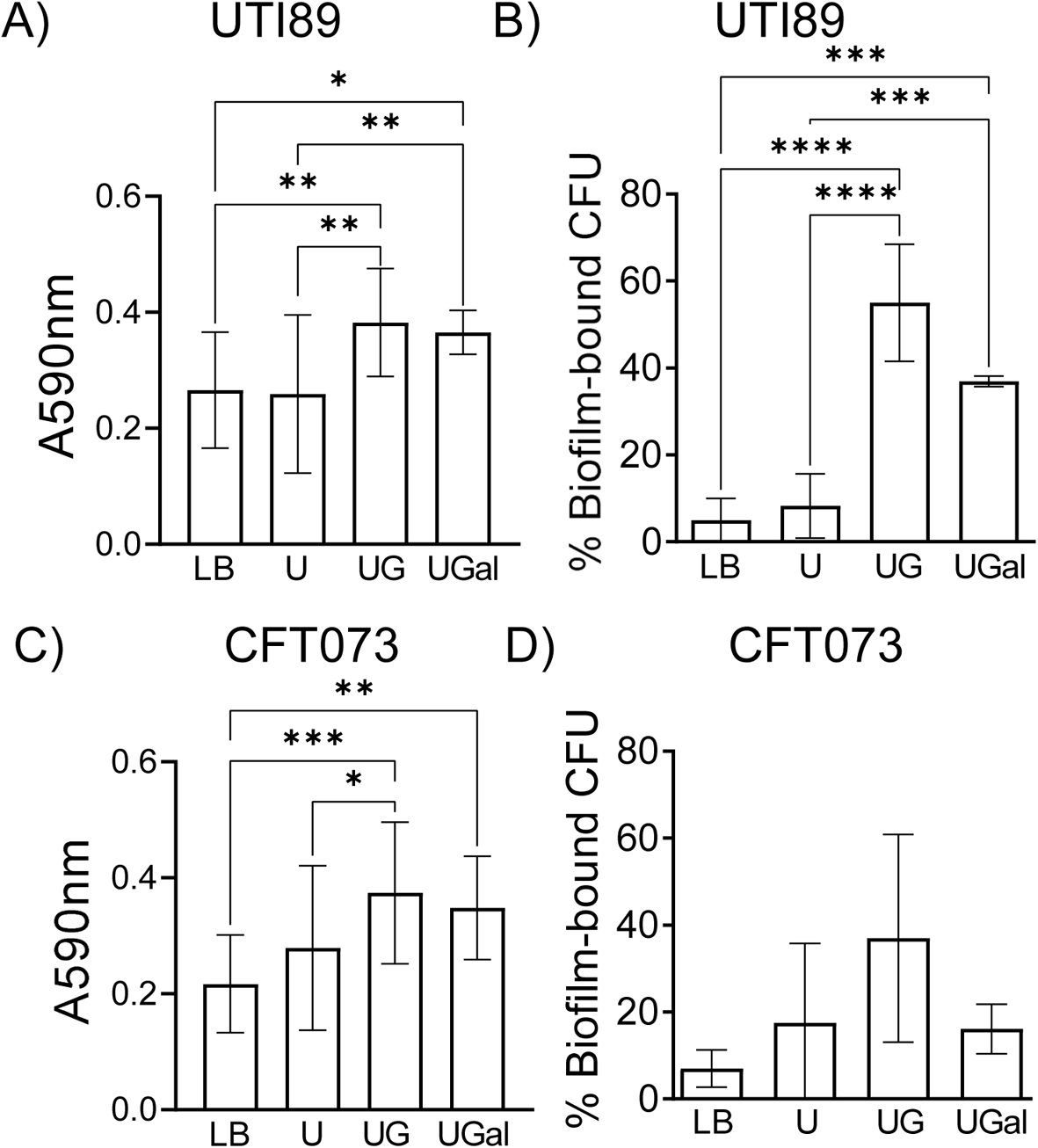
Biofilm formation by UPEC strains in the presence of human urine plain or supplemented with glucose or galactose. Type 1 piliated UTI89 (A, B) and CFT073 (C, D) were cultivated in LB, U, or UG for 24h, in 96-well polystyrene plates at 37°C, and 200rpm shaking. Biofilms were quantified by staining biomass with crystal violet and presented as histograms of average of at least four biological repeats (each with triplicate technical repeats) ± standard deviation. In separate wells, biofilm-bound and planktonic CFUs were enumerated by dilution plating and presented as average % biofilm bound CFUs from a minimum of three biological replicates ± standard deviation. Results were compared by ordinary one-way ANOVA followed by Tukey’s multiple comparisons test. *, *P*<0.05; **, *P*<0.005; ***, *P*<0.0005.

**Figure 6:**
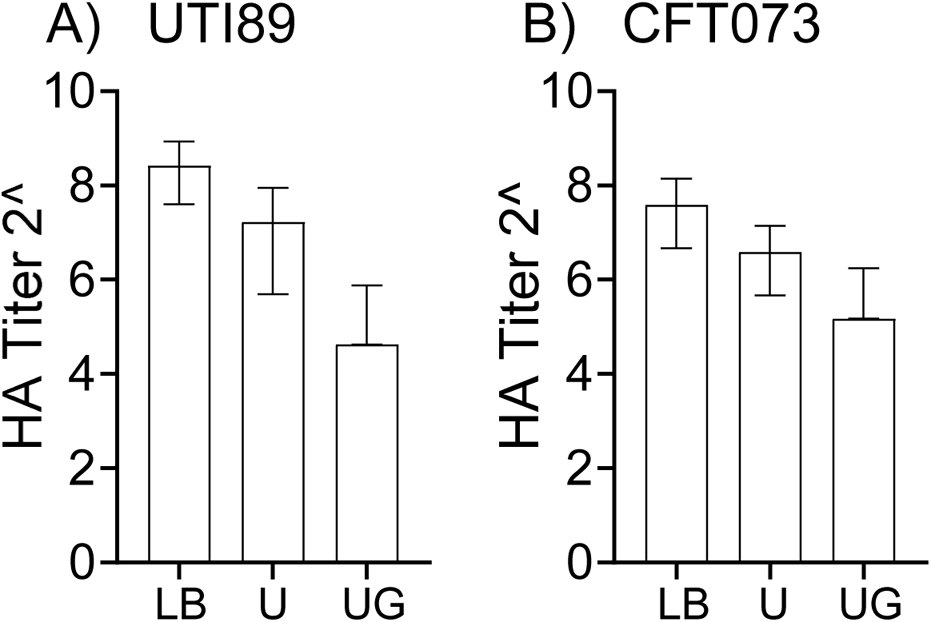
Hemagglutination (HA) titers of UPEC strains exposed to human urine ± glucose. Type 1 piliated UTI89 (A) and CFT073 (C) were exposed for 2h to LB, U, or UG washed, and adjusted to OD_600_=1.0. Bacteria were diluted two-fold (from 2^1^ to 2^10^) and mixed with Guinea pig RBCs. After incubation at 4°C for 30 minutes, hemagglutination titer was determined visually as the lowest dilution at which RBC button is visible in a v-bottom plate. Average HA titer ± standard deviation from 3 biological replicates for UTI89 and 2 biological replicates for CFT073 are shown.

**Figure 7:**
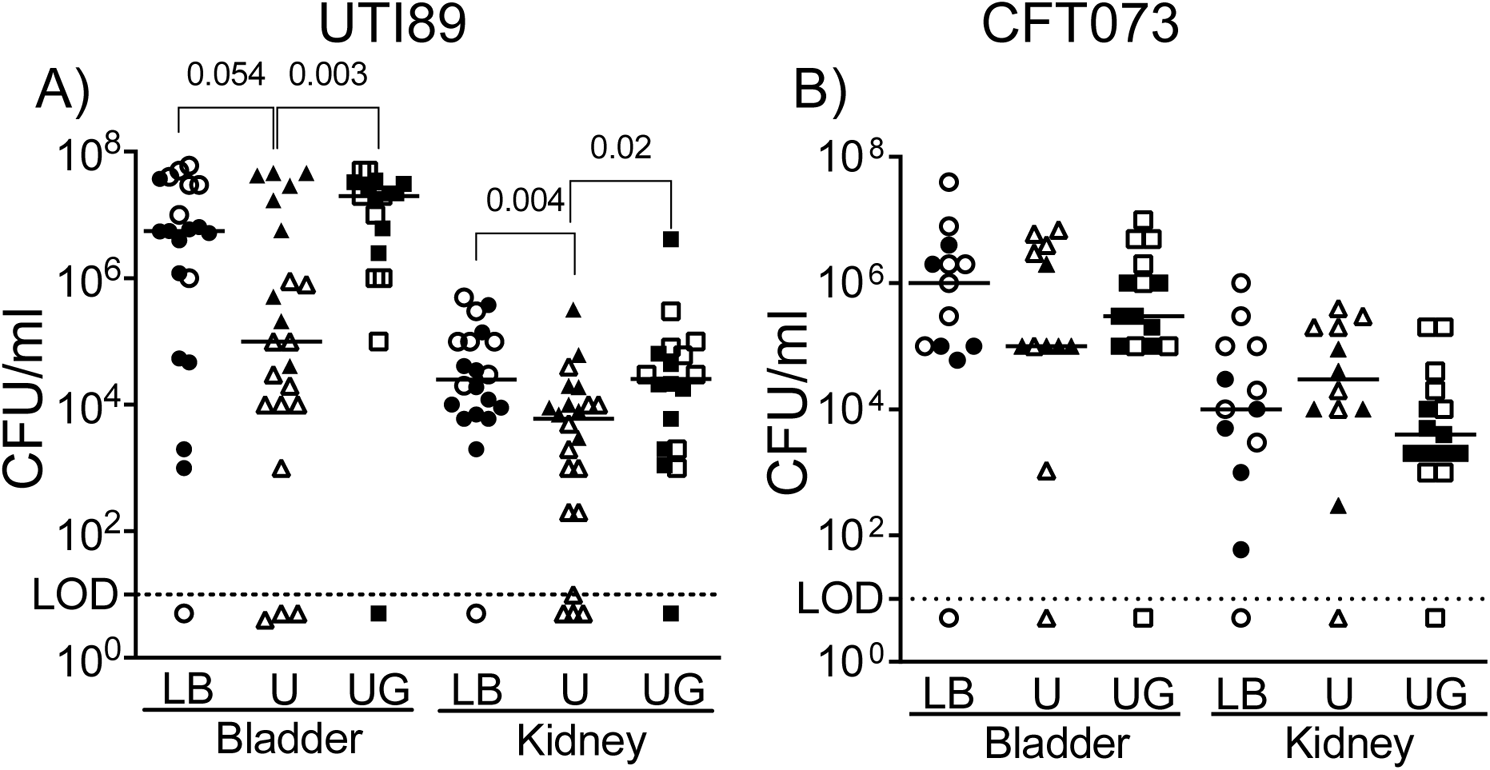
Effects of 2h-long exposure to human urine ± glucose on the uropathogenesis of UPEC strains in a mouse model of ascending UTI. Type 1 piliated UTI89 (A) and CFT073 (C) were exposed for 2h to LB, U, or UG and washed. C3H female (closed symbols) and male (open symbols) were infected with bacteria from each exposure. At 24h post-infection, bacterial organ burden in bladder and kidneys was determined by dilution plating the organ homogenates. Scatter plots show data for individual animal with median as the measure of central tendency. Data were analyzed by Mann-Whitney U test, individual P values are shown when *P*<0.05 or when *P* is very close to 0.05 (for example; *P*=0.054). Non-significant *P* values are not shown.

## DISCUSSION

A healthy urinary tract is a moderately oxygenated environment containing urine made of low concentrations of amino acids and short peptides and trace amounts of transition metals including irons (33–35). Using mouse model of ascending UTI, researchers have compared WT UPEC and deletion mutants targeting key enzymes in important metabolic pathways to establish importance of peptide transport across bacterial membrane, D-serine metabolism, acetate metabolism, gluconeogenesis, and TCA cycle in uropathogenesis of UPEC (36, 46–51). Hence, we examined UTI89-LB, UTI89-U, and UTI89-UG transcriptomes for specific metabolism genes from these pathways (Table 1–3). Compared to LB control, UTI89 exposed to either U or UG showed upregulation of genes encoding import systems for dipeptides and amino acids arginine, serine and histidine which correlates with observations from the examination of transcriptome of UPEC isolated from the mouse urinary tract (35). In addition, UPEC mutants deficient in peptide transport (*ΔdppA*) have also been observed to be impaired in colonizing murine bladder and kidney tissues; UPEC auxotrophs for arginine, serine, or glycine are also not defective in a mouse model of ascending UTI indicating that UPEC is able to import these amino acids from the urinary tract and metabolize them (49). The expression of oligopeptide transporter system genes *oppA/B/C* (UTI89_C1441/42/43) was not significantly altered in either UTI89-U or UTI89-UG, although CFT073 *ΔoppA* mutant shows impaired urinary colonization in a mouse model (49). Amino acid D-serine accumulation and catabolism to acetate supports UPEC growth and colonization in the urinary tract via regulation of expression of virulence genes encoding P, F1C fimbriae and hemolysin (36, 46–48). We observed that UTI89-U exhibits significant upregulation of D-serine catabolizing D-serine deaminase (*dsdA*, UTI89_C4955), and D-serine specific transporters (*dsdX*, UTI89_C4956), *cycA* (UTI89_C4817) and *sstT* (UTI89_C3527). Modest downregulation of *dsdA* in UTI89-UG compared to UTI89-U agrees with previous report that *dsd* operon is subject to catabolite repression in presence of glucose (52). As the nutrients in their surroundings are depleted, bacteria including UPEC activate acetate switch to utilize acetate by activating acetate-coA ligase (*acs*, UTI89_C4659), which was downregulated in UTI89-UG. These results suggest that 2h-long exposure to urine ± glucose does not activate acetate metabolism in UTI89. To determine whether UTI89 *opp* genes are activated, *dsdA* is significantly downregulated, and/or acetate metabolism is activated at time points later than 2h in an *in vivo*system, and in general to determine the importance of transport and metabolism of peptides in the pathogenesis of UPEC-UTI in diabetes, future experiments comparing WT UTI89 with mutants deficient in transport and utilization of peptides and amino acids in diabetic and non-diabetic mouse models of ascending UTI are warranted.

We observed that *fumC* was significantly induced in UTI89-U vs UTI89-LB, while expression of UTI89_C1800/C4715 encoding *fumB* class II fumarate hydratases was unaffected in UTI89-U. *fumC* was downregulated in the UTI89-UG vs UTI89-U. These results concur with previous observations that iron-restricted conditions such as those in the urinary tract favor expression of *fumC*, a class I fumarate hydratase that does not require iron for activity, and that the growth of *ΔfumC* mutant is delayed under iron-limiting conditions (51, 53). UPEC mutants *ΔsdhA* (succinate dehydrogenase) and *ΔfumC* (class II fumarate hydratase) targeting enzymes catalyzing oxidative TCA cycle are defective in a mouse model of ascending UTI, while the uropathogenesis of *ΔfrdA* (fumarate reductase) and *ΔfumB* (class I fumarate hydratase) targeting reductive TCA cycle is unaffected (49–51). Thus, oxidative TCA cycle activity is critical for uropathogenesis of UPEC, while reverse TCA cycle is dispensable for it.

The presence of glucose in urine (glycosuria) constitutes a ready source of carbon that can be converted via glycolysis to pyruvate, which can be further metabolized through either oxidative TCA cycle or to acetate or lactate. Indeed, the increased expression of genes encoding glycolytic enzymes in UTI89-UG (Table 2) suggest a switch toward glycolytic metabolism in presence of glycosuria. To address whether such metabolic switch provides a direct fitness advantage to UTI89 in diabetic urinary tract, we inoculated UTI89 and CFT073 exposed for 2h to either LB, U, or UG into male and female C3H mice via transurethral catheterization. Results show that compared to UTI89-LB control, UTI89-U is significantly defective, while UTI89-UG has modest fitness advantage in the colonization of murine bladder and kidneys. A similar trend is observed for CFT073, although differences in bladder CFU burden for CFT073-LB, CFT073-U, and CFT073-UG are statistically not significant. It is evident that compared to their UTI89-LB infected counterparts, reduction in median bladder CFUs of UTI89-U infected mice is driven primarily by the reduced bladder bacterial burden in UTI89-U-infected male mice, we have not delineated specific molecular mechanisms underlying reduced organ burden in mice infected with urine-exposed UPEC compared to those infected with UPEC-LB. Previous work in CFT073 has shown that deletion mutants targeting irreversible steps in glycolysis, *pfkA* (phosphofructokinase) and *pyk* (pyruvate kinase) are not defective in a mouse model of ascending UTI (49). We also did not detect changes in the expression of *barA/uvrY* (UTI89_C3156/C2115) two component system, which regulates a switch between glycolysis and gluconeogenesis and has been shown to be critical to the uropathogenesis of UPEC in a monkey cystitis model (54).

Catalytic activity and structural stability of almost half of enzymes from both the pathogen and its eukaryotic host are dependent on acquisition of transition metals iron, manganese, nickel, cobalt, zinc, and copper, which are critical cofactors (55). Due to the toxicity associated with their presence in free ion form, transition metals are sequestered in various host proteins such as heme, ferritin, transferrin, and ceruloplasmin; metal sequestration also give host competitive advantage over pathogens. We observed that exposure to glycosuria (UTI89_UG) rapidly and uniformly activated genes encoding free metal ion uptake systems (*sitA/B/C/D*), siderophore-mediated metal ion uptake systems (*fhu*-aerobactin, *fep*-enterobactin, *iro*-salmochelin*, ybt*-yersiniabactin), and importers of host-protein-conjugated metal ions (Chu and Hma). In contrast, 2h-long exposure to urine only activated *ybt-fyuA* yersiniabactin genes and *nikA* nickel transporter. Future experiments examining the transcriptomes of UTI89 isolated from diabetic and non-diabetic mouse bladders and kidneys at different times post-infection will help determine whether genes encoding metal acquisition systems and other virulence/regulatory factors are more rapidly induced in diabetic urinary tracts than in non-diabetic urinary tracts. Comparison of the pathogenesis of metal ion transport deficient strains in diabetic and non-diabetic mice is also needed.

We also observed that for both UPEC strains, 2h-long exposure to urine significantly reduces HA titer suggesting a corresponding reduction in assembly and/or function of type 1 pili; HA titer was further reduced in UPEC strains exposed to glycosuria. Correspondingly, in both UTI89-U and UTI89-UG (compared to UTI89-LB), we also observed significant upregulation of *fimE* (UTI89_C5010) and significant downregulation of *fimB* (UTI89_C5009), while the expression of polycistronic *fim* operon and was unaffected. FimE and FimB are trans-acting, DNA-binding proteins, of which FimE binding inverts *fimS* to phase OFF orientation and prevents transcription of *fim* operon encoding proteins involved in type 1 pilus assembly (56). Human urine not only inhibits the expression of *fim* operon by inducing *fimS* phase OFF orientation in UTI89 cultivated under type 1 piliation conditions (static, 37°C, two 24h cultures see methods), but also suppresses the function of FimH adhesive tip (57). Glucose also reduces type 1 piliation by reducing levels of cAMP—CRP in turn affecting FimB mediated inversion of *fimS* to phase ON orientation (45). However, transcript levels of *cyaA* (encodes adenylyl cyclase enzyme catalyzing cAMP synthesis, UTI89_4365) and *crp* (encodes CRP, UTI89_C3860) were unaffected in UTI89-LB, UTI89-U, and UTI89-UG. In our experiments, we inoculated plain urine or urine supplemented with glucose with type 1 piliated UPEC and maintained cultures at 37°C, static conditions for 2h-long. Thus, our results suggest that healthy urine as well as glycosuria rapidly inhibit formation and/or function type 1 pilus by UPEC strains.

Cultivation of UPEC strains in UG showed augmentation of biofilms compared to bacteria in either LB-control or plain urine. When visualized by biomass staining with crystal violet, biofilm augmentation was modest. In contrast, enumeration of biofilm-bound and planktonic CFUs by dilution plating showed a more robust augmentation of UPEC biofilms in presence of UG because biofilm-bound CFU accounted for <10% of total bacteria in both UTI89-LB and CFT073-LB. Addition of equal amount of substitute sugar, galactose to urine also significantly increased biofilm formation compared to LB-control. Thus, biofilm augmentation in UTI89-UG and CFT073-UG appears to be modulated at least partially via sugar-mediated changes in the osmolarity of urine. Both UTI89-U and UTI89-UG showed significant upregulation of *ybt-fyuA* yersiniabactin iron uptake system, which was previously shown to play a role in biofilm formation (23). Although, we did not observe reduction in biofilm formation upon addition of iron chelator to either in LB, U, or UG.

Glucose-mediated catabolite repression also inhibits biofilm formation, as glucose reduces cAMP—CRP an inducer of biosynthesis of surface adhesive type 1 pili, curli fibers, flagella, and K1 capsule (38, 39). Compared to UTI89-U, UTI89-UG showed significant downregulation of *csgD* (UTI89_C1161) and *flhC/D* (UTI89_C2094/2095), positive regulators of curli fiber and flagellar gene expression, respectively; while K1 capsule encoding operon (UTI89_C3362— UTI89_C3367) was significantly downregulated in UTI89-UG compared to UTI89-LB. As our transcriptome data show transcript levels following 2h-long exposure to LB, U, or UG it does not represent gene expression changes in UPEC strains cultivated overnight in the presence of LB or urine ± glucose for biofilm formation. Overall, to further elucidate implications of these observations to UTI pathogenesis, future experiments examining expression of *fim*E/B regulators and *fim* operon as well as experimental analysis of FimH adherence activity in UPEC from diabetic and non-diabetic murine urinary tracts are warranted.

In summary, results presented in this report provide significant insights into early changes in UPEC physiology mediated by human urine ± glucose (U or UG). For this, we cultivated UPEC in LB, a nutrient-rich medium that does not contain glucose, washed, and then exposed for 2h to either plain human urine or glycosuria. However, an important shortcoming is that our results do not inform about what might happen to UPEC physiology during long-term infection of the urinary tract of a diabetic host. This shortcoming must be addressed by future experiments analyzing the physiology (virulence and transcriptomes) of UPEC isolated from the urinary tracts (bladder and kidney tissues) of diabetic and non-diabetic mouse models at various time points after experimental induction of ascending urinary tract infection.

## SUPPLEMENTARY FIGURES

**Figure S1:** Principal-component analysis (A) and Euclidean distance between samples (B) for UTI89-LB versus UTI89-U

**Figure S2:** Principal-component analysis (A) and Euclidean distance between samples (B) for UTI89-LB versus UTI89-UG

**Figure S3:** Principal-component analysis (A) and Euclidean distance between samples (B) for UTI89-U versus UTI89-UG

**Figure S4:** Correlation between qRTPCR and RNASeq results for UTI89-LB versus UTI89-U (A) and UTI89-LB versus UTI-89-UG (B) comparisons

